# Reinstating olfactory bulb derived limbic gamma oscillations alleviates depression

**DOI:** 10.1101/2022.02.01.478683

**Authors:** Qun Li, Yuichi Takeuchi, Jiale Wang, Livia Barcsai, Lizeth K Pedraza, Gábor Kozák, Shinya Nakai, Shigeki Kato, Kazuto Kobayashi, Masahiro Ohsawa, Magor L Lőrincz, Orrin Devinsky, Gyorgy Buzsaki, Antal Berényi

## Abstract

Although the etiology of major depressive disorder remains poorly understood, impairment of gamma oscillations recently emerged as a potential biomarker for major depression. The olfactory bulb (OB) is a major source of brain wide gamma oscillations and bulbectomy is an animal model for depression. Here we demonstrate that chemogenetic suppression of OB neuronal activity or temporally suppressing the OB to pyriform cortex synaptic transmission decreased gamma oscillation power in multiple brain areas associated with depression-like behaviors. To assess the hypothesized link between depression and diffuse depression of gamma oscillations, we employed gamma phase-dependent closed loop neuromodulation of cortical areas, paced by the native OB output. This procedure alleviated depressive-like behaviors in animals and suggests that restoring gamma oscillations may improve depression in humans.

**One Sentence Summary:** Role of limbic gamma oscillations in depression

## Main Text

Major depressive disorder (MDD) is a common, severe, debilitating psychiatric illness (*1*) often resistant to pharmacotherapy (*2*). The incidence and prevalence of MDD are increasing, with COVID-19 driving more than 50 million new cases (*3*). Electroconvulsive therapy is effective in some cases but is often complicated by long-term impairments in memory and other cognitive function (*4*). Deep brain stimulation (DBS) and transcranial magnetic stimulation (TMS) are potential MDD therapies, but their long-term efficacy is uncertain (*5–8*). For drug-resistant MDD patients, alternative therapies are needed.

Coherent gamma oscillations (30-80 Hz) link brain areas by creating temporal “windows” to transfer information, characterized by enhanced excitability (*9–13*). Neuronal entrainment to gamma oscillations (*14*) and gamma coupling between limbic areas (*15–17*) can influence affect and emotional salience of stimuli. MDD characterized by neuronal network dysfunctions reflected in spectral disturbances in electroencephalographic (EEG) signals (*18*). Limbic gamma power and its long-range desynchronization are recommended MDD biomarkers (*19, 20*). Ketamine has potent antidepressant effects in humans (*21*) and animal models of depression (*22–24*), and can increase brain-wide gamma power (*25, 26*). An important physiological source of gamma oscillations is the olfactory bulb (OB) (*27*) (*28*). Nasal occlusion (*29, 30*) can suppress gamma oscillations in the primary olfactory cortex (i.e. piriform cortex, PirC) (*31*) and nucleus accumbens (NAc) (*32*). Critically, bilateral olfactory bulbectomy (OBx) rodents brings about symptoms concordant with human MDD (*33*), supporting the hypothesis that OB-driven gamma oscillations are critical for normal mood, although alternative interpretations have also been considered (*34–37*).

Gamma oscillations occur in awake naïve rats (Fig. 1B and Fig. S1 A) and are highly coherent between OB and multiple brain regions (Fig. 1B and Fig. S1 B–D). OBx dramatically reduces gamma oscillations (Fig. S1 E and F). In the PirC, the main target of OB efferents, gamma oscillations were markedly attenuated in OBx rats compared to controls (Fig. S1 G and H, for descriptive statistics, tests, and sample sizes, see Table S1 and S5). Rats developed depressive-like behaviors, including signs of anxiety (avoidance of open field, Fig. S1 I) and anhedonia (smaller sucrose water consumption, Fig. S1 J) one month following OBx, supporting that OB drives brain-wide coherent gamma oscillations are relevant to maintain a healthy mood.

**Fig. 1.**
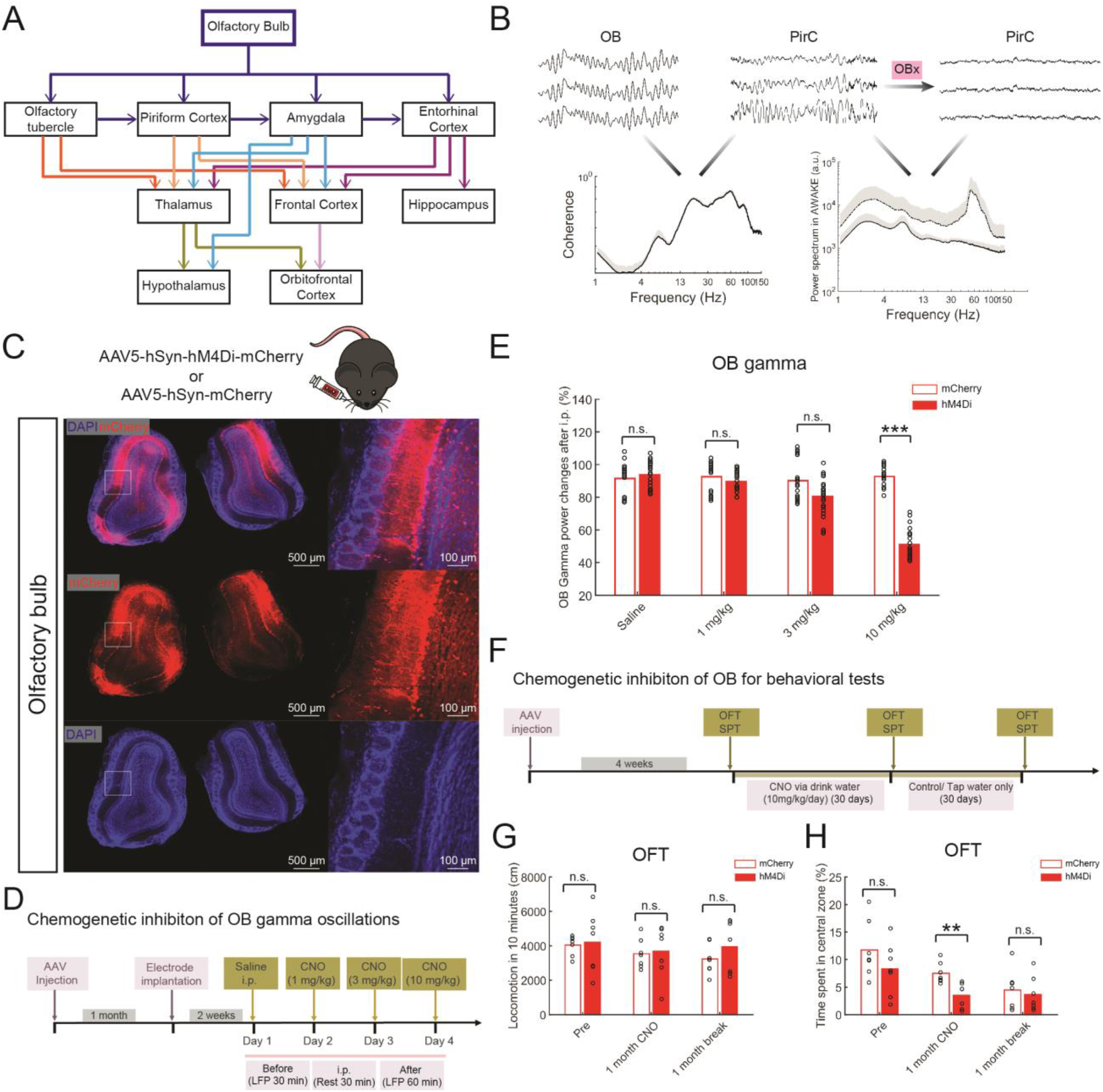
Chemogenetic inhibition of olfactory bulb neuronal activity reduces local gamma oscillations and induces depressive-like behaviors. (**A**) Major projections of olfactory bulb (OB). (**B**) Top panel, represents LFPs (0.5 s) of OB and piriform cortex (PirC) from an awake rat, and LFP in PirC after olfactory bulbectomy (OBx). Bottom panel, coherence and power spectrum corresponding to signals shown on the top panel. (**C**) Representative fluorescent images of the mouse olfactory bulb following injections of AAV5-hSyn-hM4Di-mCherry. (**D**) Schematics and timeline of the chemogenetic inhibition of OB gamma oscillations. (E) Effect of systemic administration of clozapine N-oxide (CNO) on OB gamma power of hM4Di and mCherry expressing mice, respectively (see Fig. S2 for the same protocols carried out in the rats). (**F**) Schematics and timeline of the chemogenetic inhibition of OB for behavioural tests. (**G**) Effects of CNO on the total distance travelled in the Open Field Test (OFT) for the hM4Di and mCherry expressing mice, respectively. The tests were performed before CNO administration (Pre), following 30 days of systemic CNO administration (one month CNO) and following 30 days after the cessation of the CNO treatment (one month break) (n = 7 animals/group). (**H**) Decreased time spent in the center zone during the OFT of the hM4Di group one month after CNO treatment (n = 7 / group). Results of both mean value and statistical tests are reported in detail in Table S1 and S5. n.s. indicates non-significant difference. * and ** indicate differences of P < 0.05 and P < 0.01, respectively.

To test the role of OB derived gamma oscillations in depression more directly, we generated and validated a novel oscillopathy-based (*38*) depression model lacking the confines of the ablation-based OBx model. We used Designer Receptors Exclusively Activated by Designer Drugs (DREADD)-based chemogenetic tools (*39–41*) to reversibly silence pan-neuronal activity in the OB by bilaterally injecting AAV5-hSyn-hM4Di-mCherry (Fig. 1C), a modified muscarinic acetylcholine receptor selectively activated by clozapine *N*-oxide (CNO) in the OB. After systemic CNO administration, OB gamma power (30-80 Hz) was suppressed in a dose-dependent manner in both mice (Fig. 1D and E) and rats (Fig. S2A). To evaluate the behavioral influence of long-term suppression of OB-induced gamma activity, mice were chronically treated with CNO (Fig. 1F). The hM4Di group showed anxiety-like behavior with less time spent in the center of the open field (Fig. 1H), (but no difference in overall locomotion (Fig. 1G), no significant difference in sucrose preference test (SPT) nor in daily liquid consumption (Table S6, Fig. S3A-CB). Thus, chronic silencing of OB neuronal activity suppresses OB gamma activity and facilitates depressive-like behaviors, supporting that dysfunction of OB gamma activity contributes to depression.

The role of OB to PirC projections in generating and maintaining coherent limbic gamma band activity (Fig. S1 B-D) was confirmed by pathway-specific optogenetic approaches. First, OB neurons projecting to PirC were selectively tagged by simultaneous viral vector injections of AAV5-EF1α-DIO-iC++-EYFP in OB and AAV2R-CAGGs-Cre-myc in the ipsilatral PirC, tagging the whole axonal arbors of ipsilateral PirC projecting OB neurons (Fig. S4 A). The OB to PirC was exclusively unilateral (Fig. S4 B and C), confirming previous reports (*42, 43*). We suppressed synapses with Chromophore Assisted Light Inactivation (CALI) to disrupt the OB to PirC projection by a single, brief light pulse, without affecting collateral pathways in a spatially and temporally precise manner (*44, 45*). The injection of AAVDJ-CAGGS-Flex-SYP1-miniSOG-T2A-mCherry to OB and AAV2Retro-CAGGS-Cre-myc to PirC (Fig. 2A) lead to viral expression exclusively in the PirC projecting OB neurons (Fig. 2B).

**Fig. 2.**
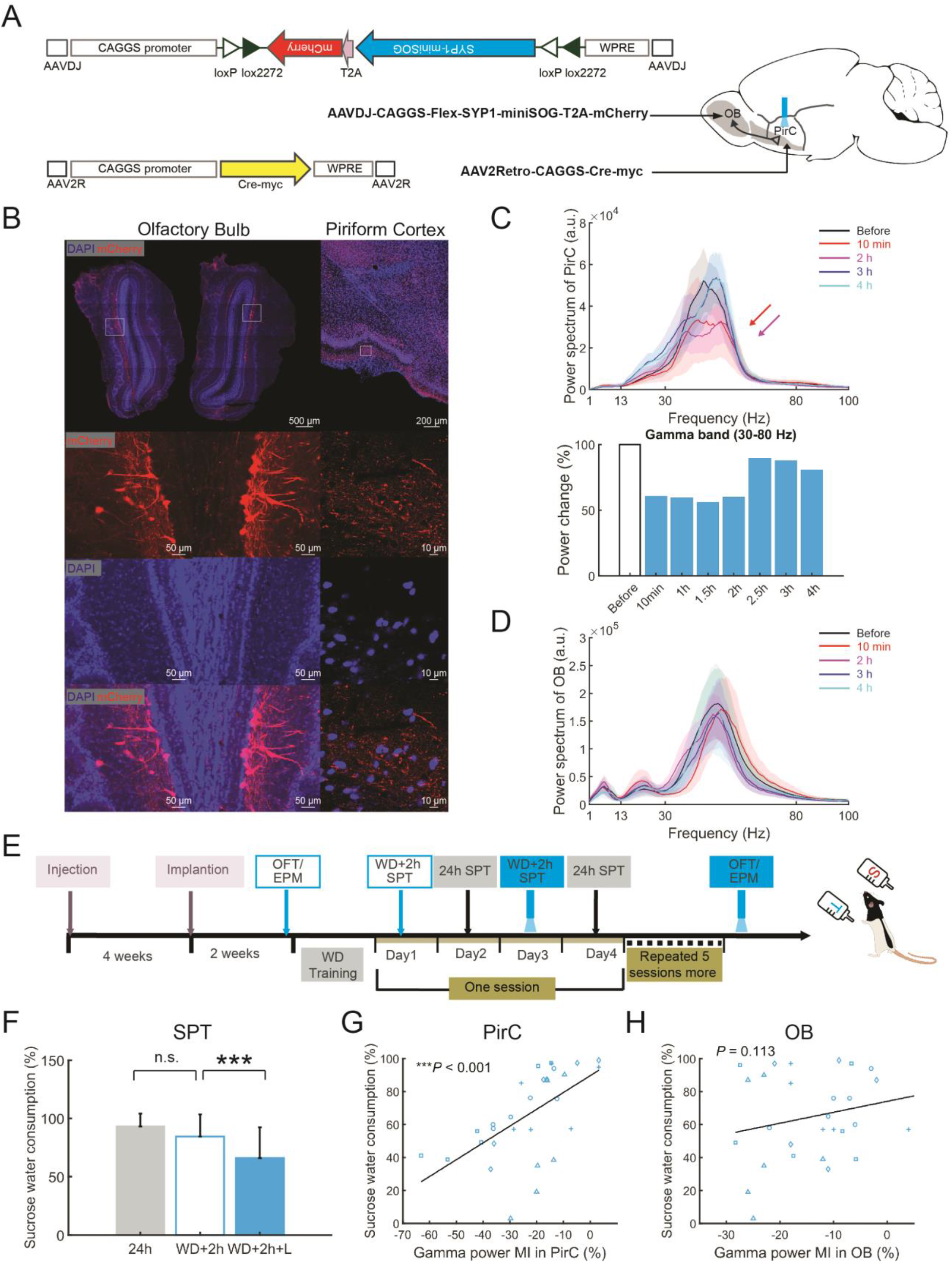
Suppressing OB to PirC synaptic transmission decreases gamma oscillations in the PirC and deteriorates performance in the sucrose preference test. (**A**) Schematics of the experiments and construct design for CALI (chromophore-assisted light inactivation) used for the specific inhibition of OB to the PirC synaptic transmission. (**B**) Fluorescent images showing mCherry expression in PirC targeting OB neurons and their axonal projections. (**C**) The suppression of gamma power in the PirC lasted for around 2 h after one time illumination (450 nm light with 20 Hz for 9 min at 9 mW at the tip). Upper panels show gamma band spectrograms (30–80 Hz) before, 1 h after (1 h) and 4 h after the illumination (4 h). Bottom panel shows quantified gamma power changes in various conditions. (**D**) Schematics of the behavioural tests following the suppression of OB to PirC synaptic transmission using miniSOG. WD: 22 h water deprivation. (**E**) Photostimulation of the PirC of miniSOG expressing rats (WD + 2 h SPT + L) decreased sucrose water consumption (120 trials from five rats). See performance of individual rats in Fig. S5A. (**F**) The disrupted sucrose preference performances were positively correlated with gamma power decrements in the PirC. Values are represented as means + S.D. Each marker represents each animal. n.s. indicates not significant difference. *** indicates difference of P < 0.001.

We characterized the effect of a single train of illumination by recording OB and PirC LFP of an awake freely moving rat before and at various post-stimulation times. Gamma oscillations were suppressed for 2h following photostimulation (PS) (Fig. 2C) in the PirC, but not in the OB (Fig. 2D). We designed an SPT accounting for these temporal constrains (Fig. 2E). Following bilateral PirC PS, the animals showed significantly lower performance in the SPT task (Fig. 2F). Further, sucrose consumption was positively correlated with gamma power in the PirC (Fig. 2G), but not OB (Fig. 2H). No significant changes were found in OFT and Elevated Plus Maze test (EPM) following PirC PS (Fig. S5 B-E). These findings suggest that short-term inhibition of OB to PirC synaptic transmissions (~2h) decreases PirC gamma oscillations resulting in anhedonia, and the magnitude of the OB derived gamma power in the PirC predicts the magnitude of depressive symptoms.

Our OB-PirC gamma phase analysis revealed a coherent phase lag, indicating an inherent oscillatory entrainment through direct synaptic connection (*46*) (Fig. S6). We investigated whether a long-term entrainment by OB-derived PirC gamma oscillations affects depressive behaviors. We developed an unsupervised real-time closed-loop intervention to modify PirC gamma oscillations via phase-locked (InPhase, AntiPhase) electrical stimulation driven by the detected gamma oscillations in the OB (Fig. 3A-C). AntiPhase gamma E-Stim (i.e. interfering with PirC rhythmic neuronal activity) decreased sucrose preference in all animals and the effect outlasted the stimulation (Fig. 3D and Fig. S7). In contrast, InPhase stimulation had no effect on sucrose preference (Fig. 3D). Neither InPhase nor AntiPhase stimulation affected the rats’ spontaneous locomotion in their homecage (Fig. 3E). AntiPhase stimulation decreased gamma power in the PirC while InPhase E-Stim increased gamma power (Fig. 3F, G and Fig. S8 A). Neither InPhase nor AntiPhase changed gamma frequency distribution in the PirC (Fig. S8 B) and the incidence of gamma events during the awake state was unaltered (Fig. 3H). These results suggest that closed-loop OB gamma neuromodulation of PirC enhances and silences PirC gamma oscillations in a phase-dependent manner and the effects persist for at least one day after stimulation. The AntiPhase gamma stimulation and resulting decrease in PirC gamma oscillations caused anhedonia, consistent with a depressive-like state.

**Fig. 3.**
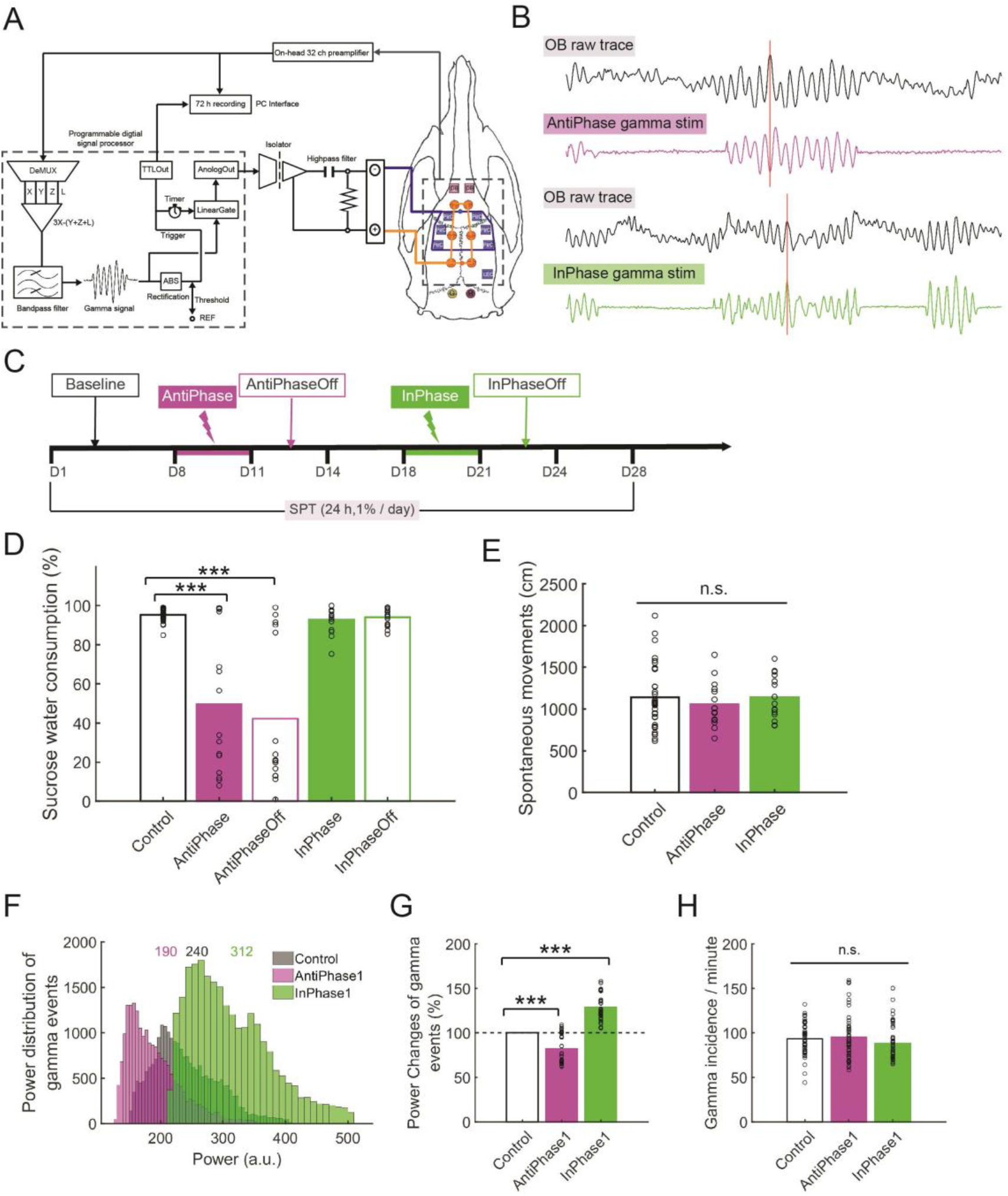
Real-time silencing of OB-origin gamma oscillations to the PirC phase-dependently induces anhedonia in naïve rats. (**A**) The schema of closed-loop neuromodulation of the PirC with OB-origin gamma oscillations in real-time. The pre-amplified and multiplexed LFP signals were fed to a programmable digital signal processor. The signals were demultiplexed and analized online to gamma events in the OB using a custom-made signal detection algorithm. LFP signals were demultiplexed at 500 Hz per channel and a signal from a pre-selected OB channel was band-pass filtered with a 4th order Butterworth filter to 30–110 Hz. Common artifacts were removed from the selected channel X (from OB) by subtracting averaged signals in the left (Y) and right (Z) PirC, and LEC (L). Each 300 ms-electrical stimulation in filtered OB gamma wave signal was triggered when a filtered signal exceeded a fine-tuned arbitrary threshold for each animal. Time resolution of the detection was 2 ms. The gated gamma E-Stim signal was fed to the six PirC locations via an analog isolator IC (ISO124, Texas Instruments) through a RC high-pass filter of 0.25 s time constant with a present phase delay. Orange circles represent miniature machine screws as cathodal (i.e. returning) electrodes. Squares indicate temporal cranial windows through which the recording and stimulating electrodes were introduced. Blue ones were for the PirC and LEC, while purple ones were for OB. (**B**) Representative LFP raw traces of OB and three derivatives for different phase stimulation (AntiPhase, InPhase). Detected OB gamma oscillations were fed to the PirC with AntiPhase and InPhase respectively. Red vertical lines indicate positive peaks of original gamma oscillations in the OB. ‘Upward’ signal represent neuronal activity (following the EEG polarity conventions) and cathodal current on the OB and stimulus traces, respectively. (**C**) The schema of the experiment. Each stimulation was carried out for three days continuously, followed by an additional three days without stimulation (OFF days). (**D**) AntiPhase stimulation significantly decreased sucrose water preference in the naïve rats, and the effects lasted for 1-4 days even after termination of the stimulus. See also induvial five trials in the Fig. S7. InPhase didn’t decrease sucrose water consumption. (**E**) No significant side-effects of the E-Stim on spontaneous movements in the homecage was observed. (**F**) The sum of power distribution of gamma events in the PirC on the day before the first stimulation (Base), during the first day after AntiPhase stimulation ended (AntiPhase1) and during the first day after InPhase stimulation ended (InPhase1). The numbers represent medians of the distributions. See also individual five trials in the Fig. S8. (**G**) AntiPhase stimulation significantly decreased power of gamma events in the PirC whereas InPhase stimulation increased it. (**H**) No significant differences of incidence of gamma events between Base, AntiPhase1 and InPhase1 in the awake state. Values are represented as means with bars. Each circle represents each trial. n.s. indicates not significant difference. * and *** indicate difference of P < 0.05 and P < 0.001, respectively.

To investigate whether boosting PirC gamma oscillations can alleviate depressive-like behaviors, we used InPhase closed-loop PirC E-Stim in several depression models, including Lipopolysaccharide (LPS) anhedonia test in SPT, despair-like behavior in FST, anxiety-like behavior in OFT and EPM in both rats (*47*) and mice (*48*) (Fig. 4A). In the SPT, LPS induced decreased sucrose preference, but the group receiving InPhase gamma E-Stim recovered SPT performance (Fig. 4B and Fig. S9). InPhase E-Stim also increased the ‘center time’ during the OFT (Fig. 4C), number of center entries (Fig. 4D) and total distance travelled per time unit (Fig. 4E) compared to non-stimulated animals. Similarly, InPhase gamma E-Stim alleviated anxiety-like behaviors in the EPM test (Fig. 4F), and increased the total distance travelled per time unit (Fig. 4I), but did not alter the time spent in the closed arms (Fig. 4G) or in the center (Fig. 4H). In contrast, AntiPhase E-Stim failed to improve rat behavior in the OFT (Fig. 4C–E) and EPM tests (Fig. 4G–I). These results suggest that the OB derived phase-matched closed-loop gamma neuromodulation in the PirC alleviates depressive-like and anxiety-like behaviors in the SPT, OFT and EPM tests, standard rodent models of depression.

**Fig. 4.**
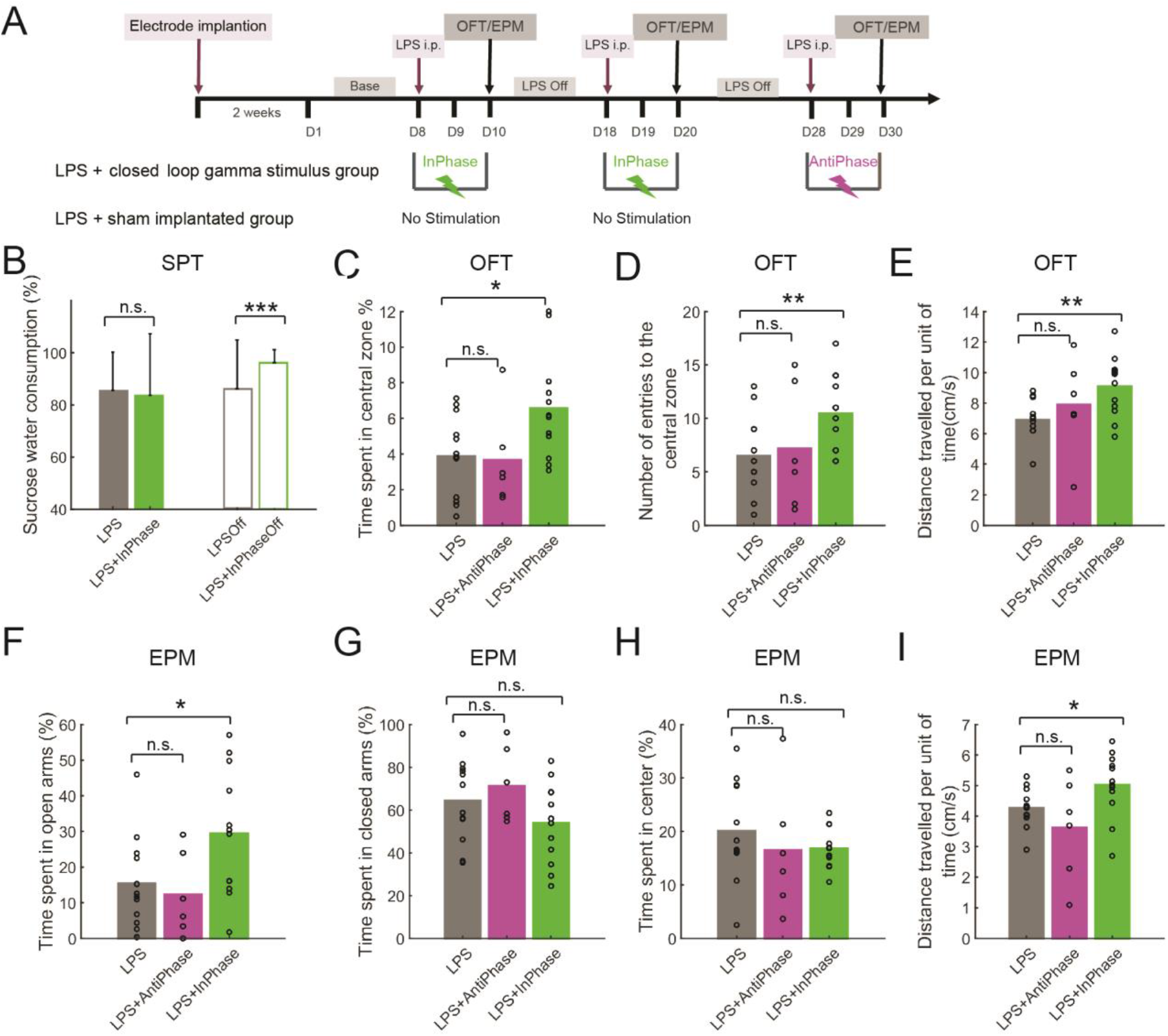
Reinstating OB-derived gamma oscillations to the PirC phase-dependently alleviates depressive-like behaviors. (**A**) Schematics of the experiment. (**B**) LPS decreased the sucrose preference in both groups by systemic administration of lipopolysaccharide (LPS). Two days of InPhase stimulation recovered the decreased SPT performance in the post stimulation days. See also performances of individual rats during the two InPhase stimulation sessions in Fig. S9 (n = 6 rats / group). (**C–I**) InPhase stimulation alleviated the depressive-like behaviors in the OFT (**C–E**) and the anxiety-like behaviours in the EPM test (**F–I**) induced by the LPS administration. Note, that AntiPhase stimulation failed to reproduce these behavioral benefits. Values are presented as means ± S.D. Circles indicate individual trials. n.s. indicates not significant difference. *, **, and *** indicate difference of P < 0.05, P < 0.01, and P < 0.001, respectively.

Our experiments reveal a mechanism between deficient gamma activity and behavioral decline in the OBx model of depression. The OBx model includes neurochemical, neuroanatomical, physiological, endocrine and behavioral alterations concordant with MDD (*49*). The brain-wide suppression of gamma oscillations in OBx animals and gamma oscillopathies in MDD patients share several resemblances (*19, 50*). Further research should strengthen the link between the bidirectional manipulation of OB-derived gamma oscillations in the PirC and the corresponding behavioral changes, but the importance of PirC in network interactions underlying depression is strongly supported (*51–53*). The PirC is connected to most emotional limbic regions (*54, 55*). The dynamics of neuronal activity in multiple brain regions can affect emotional states (*56–58*) and manipulations within one circuit node can alter activity across brain regions and influence emotional behavior (*53, 59, 60*). Thus, gamma oscillations originating from the OB can alter emotional behavior through the PirC and its widespread limbic connections. Gamma oscillation can efficiently organize assembly dynamics within cortical networks (*13, 61*) and reactivating cellular ensembles in the limbic system, primed during positive experience, can ameliorate depression-like behaviors (*62*). In addition to specific sensory responses, PirC neurons respond to reward, noxious stimuli (*63*) and encode multimodal, hedonic, and context-dependent components of the representation (*64*). PirC stimulation sustains self-stimulation, presumably by inducing an internally rewarding state (*65*). Our results, demonstrating the power of closed-loop gamma-enhancement, offer a potential therapeutic route for alleviating depression-related behaviors.

## Acknowledgments

We thank Dr. Péter Hegyi for providing access to confocal microscopy. We also thank Karl Deisseroth, Bryan Roth, and Roger Tsien for their gifts of pAAV-hSyn-mCherry (Addgene #114472), pAAV-hSyn-hM4D(Gi)-mCherry (Addgene #50475), and pAAV-SYP1-miniSOG-T2A-mCherry (Addgene # 50972), respectively.

## Author contributions

Conceptualization: AB

Methodology: QL, YT, GB, AB

Investigation: QL, YT, AB

Visualization: QL, JW, LB, LP

Software: QL, YT, GK

Formal Analysis: QL

Resources: YT, SN, SK, KK, MO, AB

Funding acquisition: YT, AB Supervision: YT, OD, GB, AB Writing – original draft: QL

Writing – review & editing: YT, MLL, OD, GB, AB

## Data and materials availability

Data included and software used in this article will be available upon request from the corresponding author

## Supplementary Materials

Materials and Methods

Fig. S1 to S9

Table S1 to S6

References 66-77

## MATERIALS AND METHODS

### Animals

A total number of 20 adult male wild-type C57BL/6 mice (3 months old, 20–30 g) and 36 wild-type male Long-Evens rats (3–4 months old, 300–400 g) were used in this study. Both rats and mice were provided with a commercial diet and water *ad libitum* under a 12 h light/dark environment (light onset at 7 A.M.). The animals were housed as groups (3 animals/cage) before surgery, and then individually for the duration of the whole experiment. All animal studies and experimental procedures were approved by the Ethical Committee for Animal Research at the Albert Szent-Györgyi Medical and Pharmaceutical Center of the University of Szeged and the Government Office of Csongrád-Csanád County, Hungary (XIV/218/2016 and XIV/824/2021) and conformed to European Union guidelines (2003/65/CE) and the National Institutes of Health Guidelines for the Care and Use of Animals for Experimental Procedures.

## Method details

### Olfactory bulbectomy experiments

#### Olfactory bulbectomy surgery

Animals were anaesthetized with 4% isoflurane initially and then mounted in a stereotaxic apparatus with 1–2% isoflurane during the whole surgery. After exposing the skull, two holes were drilled with coordinates (AP: 7 mm anterior from bregma, ML: ± 2 mm from the middle line). Bilateral olfactory bulbs were removed by suction, then holes were filled with haemostatic sponge (*66*). Control animals underwent similar surgical procedures but the olfactory bulbs were left intact. The behavioural tests were performed one month after recovery (Fig. S1 I and J).

#### Chronic implantation of recording electrodes

Tripolar tungsten electrodes for intracortical recording were prepared as previous described (*67*). For the purpose of exploring the relationship between olfactory bulb and putatively relevant brain areas in intact animals (Fig. S1 A–D), electrodes were implanted in two naïve rats into the following brain areas (see also Table S2):

• OB (olfactory bulb; AP: 8.0 mm anterior from the bregma; ML: 1.0 mm, DV: 1.4, 1.8 and 2.2 mm from the dura),
• PrL/IL (prelimbic cortex/ infralimbic cortex; AP: 3.25 mm anterior from the bregma; ML: 0.5 mm, DV: 2.0, 3.0 and 4.0 mm from the dura),
• NAc (nucleus accumbens; AP: 2.0 mm anterior from the bregma; ML: 1.5 mm, DV: 6.5, 7.0 and 7.5 mm from the dura),
• PirC (piriform cortex; AP: 2.0 mm anterior from the bregma; ML: 4.0 mm, DV: 6.5, 7.0 and 7.5 mm from the dura),
• vHip (ventral hippocampus; penetrated from the 8.3 mm posterior from the transverse sinus at a 18° caudally tilted angle from the coronal plane; ML: 4.0 and 5.0 mm, Distance: 7.0, 7.5 and 8.0 mm from the dura),
• CeA/BLA (central amygdala/basal amygdala; penetrated from the 2.2 mm posterior from the transverse sinus at a 6° caudally tilted angle from the parasagittal plane; ML: 3.0 and 4.5 mm, Distance: 7.5, 8.0 and 8.5 mm from the dura), and VTA (ventral tegmental area; penetrated from the 5.30 mm posterior from the transverse sinus at a 6° caudally tilted angle from the coronal plane; ML: 1.0 mm, Distance: 7.2, 7.6 and 8.0 mm from the dura).

To investigate LFPs in the OBx animals (Fig. S1 E–H), one animal per group was implanted at 30 recording sites after the behavioural tests. Electrodes were distributed to the right hemisphere with brain areas as follows:

• PrL/IL (prelimbic cortex/ infralimbic cortex; AP: 3.25 mm anterior from the bregma; ML: 0.5 mm, DV: 2.0, 3.0 and 4.0 mm from the dura),
• M2 (secondary motor cortex; AP: 4.2 mm anterior from the bregma; ML: 1.75 mm, DV: 1.0, 1.5 and 2.0 mm from the dura),
• mid NAc (middle nucleus accumbens; AP: 2.0 mm anterior from the bregma; ML: 1.0 mm, DV: 6.0, 6.5 and 7.0 mm from the dura),
• lat NAc (lateral nucleus accumbens; AP: 2.0 mm anterior from the bregma; ML: 2.5 mm, DV: 6.0, 6.5 and 7.0 mm from the dura),
• PirC (piriform cortex; AP: 2.0 mm anterior from the bregma; ML: 4.0 mm, DV: 6.5, 7.0 and 7.5 mm from the dura),
• ant CgC (anterior cingulate cortex; AP: 0.48 mm anterior from the bregma; ML: 0.5 mm, DV: 1.0, 1.5 and 2.0 mm from the dura),
• post CgC (posterior cingulate cortex; AP: 0.84 mm posterior from the bregma; ML: 0.5 mm, DV: 1.0, 1.5 and 2.0 mm from the dura),
• S1 (somatosensory cortex, AP: 1.20 mm posterior from the bregma; ML: 3.0 mm, DV: 1.0, 1.5 and 2.0 mm from the dura),
• EP (entopeduncular nucleus, AP: 2.4 mm posterior from the bregma; ML: 2.75 mm, DV: 7.0, 7.4 and 7.8 mm from the dura), and
• VPL (ventral posterolateral thalamic nucleus; AP: 2.4 mm posterior from the bregma; ML: 3.25 mm, DV: 5.0, 5.5 and 6.0 mm from the dura).

Recordings commenced after a recovery period of at least 14 days following surgery in the present study.

### Chemogenetic inhibition of OB neurons

For exploring gamma power changes following OB chemogenetic silencing, six mice: AAV5-hSyn-mCherry (n = 3, control), AAV5-hSyn-hM4Di-mCherry (n = 3, treated), total injection sites: 18/animal, 0.2 μl/site; and six rats: AAV5-hSyn-mCherry (n = 2, control), AAV5-hSyn-hM4Di-mCherry (n = 4, treated), total injection sites: 30/animal, 0.3 μl/site were injected (injection coordinates are shown in Table S3). One month later, electrodes were bilaterally implanted into all animals in OB (Mouse, AP: 4.8 mm anterior from the bregma; ML: ± 0.5 mm, DV: 1.4 mm from the dura; Rat, AP: 8.0 mm anterior from the bregma; ML: ± 1.0 mm, DV: 1.4, 1.8 and 2.2 mm from the dura) and PirC (Mouse, AP: 1.78 mm anterior from the bregma; ML: ± 2.0 mm, DV: 4 mm from the dura; Rat, AP: penetrated from the 2.0 mm anterior from the bregma; ML: ± 2.6 from the midline at a 10° medially tilted angle from the parasagittal plane; Distance: 6.8, 7.1 and 7.4 mm from the dura). The electrode in each area for mouse was consisted of single tungsten wire, while for rat tripolar tungsten wires were used. To test the acute modulation of OB gamma oscillations by hM4Di DREADD receptors, two weeks after surgery, saline, 1 mg/kg CNO, 3 mg/kg CNO and 10 mg/kg CNO were administrated intraperitoneally into each animal, testing one concentration per day (Fig. 1C). LFP recordings in freely-moving condition (*68*) were acquired (20 k Sample/s per channel for mouse, 500 Sample/s per channel for rat) with KJE–1001 Amplipex (*69*). In each day, we first recorded LFP from the freely-moving animals for 30 min as baseline. Then saline or CNO solutions were administered as mentioned above under five minutes of light isoflurane anaesthesia to minimize fear and pain. After 30 minutes of recovery from anesthesia we recorded post-injection LFP for 60 min. The protocol was repeated twice on each rat in two weeks.

For the chronic modulation of OB gamma oscillations by hM4Di DREADD receptors, 14 mice were injected with either AAV5-hSyn-mCherry or AAV5-hSyn-hM4Di-mCherry under 0.8–1% isoflurane anaesthetized state, respectively, by means of a standard intracranial virus vector injection technique (*70*). In total 18 injection sites (0.2 μl at each site) were distributed in the OB, details can be seen in Table S3. After at least four weeks of virus expression period, both groups received 5 ml CNO solution (10 mg/kg/day) for 30 days as drink supplement. The solution was freshly prepared every day at 7 p.m. during one month treatment. The bottles were covered with aluminium foil to avoid photolytic effects on CNO stability. Behaviour of each mouse was tested three times immediately before CNO treatment, after one month of continous CNO treatment and after 1 month of recovery period after seizing CNO treatment as shown in Fig. 1F.

### Optogenetic inhibition of the OB to PirC synaptic transmission

#### Preparation of a retrograde Cre-expresssing AAV vector

AAV vector serotype 2-R (retrograde) was prepared based on AAV Helper-Free system (Agilent Technologies), as described in Kato et al., (2018) (*71*). The transfer plasmid contained the cDNA encoding Cre recombinase and myc tag sequences downstream of the CAGGS promoter. HEK293T cells were transfected with the transfer, an adeno-helper, and expressing the adenoviral genes required for AAV replication and encapsidation plasmid through the calcium phosphate precipitation method. The crude viral lysate was purified with two rounds of CsCl gradient centrifugation, dialyzed, and concentrated with an Amicon filter (Merck Millipore). The viral genome titer was determined by quantitative PCR.

#### Preparation of an anterograde Cre-dependent miniSOG-expressing AAV vector

The core concept of the miniSOG is based on the optogenetic inhibition of synaptic release with chromophore-assisted light inactivation technique, which is fused miniSOG to the C terminus of the SYP1, and after light illumination, singlet oxygen is generated by miniSOG leading to the inactivation of fusion protein (*44*). AAV vector serotype DJ was prepared based on AAV Helper-Free system (VPK-400-DJ, Cell Biolab). The transfer plasmid contained the cDNA encoding a FLEXed InSynC (SYP1-miniSOG-T2A-mCherry) sequence downstream of the CAGGS promoter (*44*). HEK293T cells were transfected with the transfer, an adeno-helper, and expressing the adenoviral genes required for AAV replication and encapsidation plasmid through the polyethylenimimine method. The crude viral lysate was purified using discontinuous iodixanol gradients (*72*). The viral genome titer was determined by quantitative PCR.

#### Construction of optoprobes

Optical fiber was made as previously reported (*70*). A 0.39 NA, Ø200 μm core multimode optical fiber (FT200EMT, Thorlabs) was terminated with a stainless-steel ferrule (SF230, Thorlabs) and then polished in one side; exposed 1 cm the silica core, removed TECS cladding and shaped with hydrofluoric acid in another side. Make sure the selected optoprobes with maximal current output power at ~9 mW in the tip examining by a photodiode power sensor (S130C, Thorlabs) and a power meter (PM200, Thorlabs). After that, each optical fiber was glued to a single tungsten wire by a UV-curing optical adhesive (NOA61, Thorlabs), the tip of wire was 0.5 mm longer than the tip of optical fiber.

#### Behavioural and electrophysiological tests

Five adult male rats were anaesthetized with 1–2 % isoflurane and injected bilaterally AAVDJ-CAGGS-Flex-SYP1-miniSOG-T2A-mCherry in total 30 sites across the OB, and AAV2R-CAGGS-Cre-myc in total 6 sites across the PirC. The injection coordinates are presented in Table S3. Four weeks later, each rat was implanted with two tripolar electrodes in the bilateral OB (AP: 8.0 mm anterior from the bregma; ML: 1.0 mm, DV: 1.4, 1.8 and 2.2 mm from the dura) and two optoprobes in the bilateral PirC (AP: penetrated from the 2.0 mm anterior from the bregma and ML: ± 3.3 from the midline at a 5° medially tilted angle from the parasagittal plane; Distance: 6.6, 6.9 and 7.2 mm from the dura). After two weeks of recovery animals underwent water deprivation (WD) training as follows. It was consisted alternating days of i) 24 h sucrose preference test (SPT) with ad libitum access to both water and sucrose solution and ii) 22h of water deprivation followed by the sucrose preference test for 2h. Sucrose preference test was detailed as below in behavioural test part. The WD training was lasting around two weeks until all animals reached ~ 90% sucrose consumption during the 2 h SPT following WD. During the formal experiment (Fig. 2E), each session (lasting four days) contained the following blocks: i) 2 h SPT following 22 h WD, ii) 24 h SPT without WD, iii) 2 h SPT following 22 h WD, preceded by photostimulation (a 9 min long 20 Hz train of 450 nm 25 ms light pulses, 9 mW at both fiber tips in bilateral PirC) and iv) 24 h SPT without WD. To avoid building a place preference the location of the tap water bottle and sucrose water bottle were daily intermingled. These four-day long sessions were repeated six times. Open field test (OFT) and elevated plus maze (EPM) test were implemented before WD training and also after the sixth session. The second OFT and EPM tests were performed following the same photostimulation protocol as the one during the 6 test sessions previously described. For all photostimulation experiments, 10 min LFP was recorded from the PirC before and after light delivery.

### Closed-loop OB derived gamma neuromodulation of PirC

#### Electrode implantation surgery

Each animal was chronically implanted with intracortical recording and stimulating electrodes in following areas: two tripolar electrodes in bilateral OB for recording gamma oscillations for online detection, one tripolar electrode in LEC for removing global EMG noise during online gamma detection, six bipolar, combined recording and stimulating electrodes in six bilateral locations of PirC for injecting gamma activities which were detected from OB under selective modulating parameters (InPhase and AntiPhase). Bipolar stimulation electrodes in the PirC were prepared as follows, two tungsten wires axially spaced 0.3 mm apart, the tip of deep one was stripped around 0.2–0.3 mm for electrical stimulation, short wire was connected to a signal multiplexing headstage (HS3_v1.3, Amplipex, Szeged, Hungary) as same as the OB and LEC electrodes for long-term freely-moving recording’s purpose. During the chronic implantation, all six stimulus wires from six locations of PirC were combined together to one bin of connector as anode current input, and six stainless-steel machine screws were decentralized installed in the skull and then combined to the other bin of the connector as cathode (Fig. 3A). The connector physically affixed in the edge of copper mesh with a dental cement in the end of the surgery (Unifast Trad, GC). Two miniature machine screws were installed above the cerebellum as reference and ground respectively. The stereotaxic coordinates were as follows:

• bilateral OB (AP: 8.0 mm anterior from the bregma; ML: ± 1.0 mm, DV: 1.4, 1.8 and 2.2 mm from the dura),
• bilateral PirCA (anterior piriform cortex, AP: penetrated from the 2.0 mm anterior from the bregma and ML: ± 2.6 from the midline at a 10° medially tilted angle from the parasagittal plane; Distance: 6.8, 7.1 and 7.4 mm from the dura),
• bilateral PirCM (middle piriform cortex, AP: penetrated from the 0.0 mm from the bregma and ML: ± 3.5 mm from the midline at a 10° medially tilted angle from the parasagittal plane; Distance: 7.7, 8.0 and 8.3 mm from the dura),
• bilateral PirCP (posterior piriform cortex, AP: penetrated from the 2.0 mm posterior from the bregma and ML: ± 4.0 mm from the midline at a 10° medially tilted angle from the parasagittal plane; Distance: 8.4, 8.7 and 9.0 mm from the dura), and
• right LEC (lateral entorhinal cortex, AP: penetrated from 6.0 mm posterior from the bregma and 4.0 mm rightward from the midline at a 20° leftward tilted angle from the parasagittal plane; Distance: 7.8, 8.1 and 8.4 mm from the dura).

#### Long-term closed-loop OB derived gamma neuromodulation in freely-moving animals

Each stimulation block lasted for three continuous days performed in the home cage of the animals starting at 7 P.M. For achieving undisrupted online stimulation, care was taken to avoid twisting and over-tension of the cables as previously described (*68*). Briefly, a thin and light recording cable (40 AWG Nylon Kerrigan-Lewis Litz wire, Alpha Wire, Elizabeth, NJ, USA) was connected to a suspended commutator (Adafruit, New York, NY, USA) sliding vertically on guide rails to avoid the twisting and over-tension of the cables. The continuously recorded LFP signals in the OB were used to feed the endogenous gamma band oscillations by real-time closed-loop electrical stimulation to multiple sites in the PirC. The pre-amplified and multiplexed analogue LFP signals were fed to a programmable digital signal processor (RX-8, Tucker-Davis Technologies, Alachua, FL, USA) and to the data acquisition system (KJE-1001, Amplipex), both sampled at 16 kHz. The signals were demultiplexed and the OB channels were analysed online to detect gamma events using a custom-made signal detection algorithm based on a previously established routine (Kozák and Berényi, 2017). Briefly, LFP signals were demultiplexed at 500 Hz per channel and a signal from a pre-selected OB channel was band-pass filtered with a 4th order Butterworth filter to 30– 110 Hz. Common artifacts were removed from the selected channel by subtracting averaged signals in the left and right PirC, and LEC. Electrical stimulation (maximum duration: 300 ms) was triggered when a filtered artefact attenuated OB signal exceeded a fine-tuned adaptive threshold for each animal. Time resolution of the detection was 2 ms. The same filtered OB gamma signals was used as electrical stimulation (E-Stim) and when a gamma detection occurred, it was fed to the six PirC locations via an analog isolator IC (ISO124, Texas Instruments) through a RC high-pass filter of 0.25 s time constant (Fig. 3A). For InPhase stimulation, the filtered and gated gamma waveforms were fed as they were. For AntiPhase stimulation, the signals were inverted (Fig. 3B).

#### Experimental procedures for naïve animals

Two weeks after implantation, 24 h SPT was recorded every day until completed the experiment (Fig. 3 C and D). The SPT details were as following description in the Behavioural tests part. Spontaneous movements in the homecages were captured by a camera from 7–9 p.m. during one week, as baseline, and on each stimulation days (Fig. 3 E). LFP recordings were collected between 7–9 p.m. before stimulation as baseline and on the day after each E-Stim session (InPhase1/AntiPhase1, Fig. 3 F–H). Four rats were exposed to the closed-loop gamma stimulation, each receiving both InPhase and AntiPhase stimulations according to the procedure shown in Fig. 3C and Fig. S7 B (Baseline (7 days) – AntiPhase (3 days) – Off (7 days) – InPhase (3 days) – Off (7 days). The first three days of each seven day Off periods following the stimulation blocks were investigated to reveal any lasting effects of gamma E-Stim. To confirm the lack of eventual accumulating E-Stim effects from one stimulation block to another, one of the rats was also presented with an opposite order of phase sequences as follows: Baseline (7 days) – InPhase (3 days) – Off (7 days) – AntiPhase (3 days) – Off (7 days) (Fig. S7 C).

#### Experimental procedures for depression model animals

Six rats were implanted the same way as those above for closed-loop gamma E-Stim. Another six sham operated rats experienced the same surgery as the implanted group including the on-head Faraday cage made of copper mesh, but without any wire electrodes implanted. After two weeks recovery, 24 h SPT test was performed in both groups for one week as baseline. Then lipopolysaccharides (LPS, O111:B4, Sigma) dissolved in sterile 0.9% saline was injected i.p. into both groups at a dose of 200 μg/kg in a volume of 2 ml/kg, and the 24 SPT test was continued every day until the end of the experiment. After LPS injection, animals from the implanted group were immediately connected to recording/stimulating apparatus for real time InPhase gamma stimulation from 7 p.m. until the third day morning. The sham group received no stimulation. On the third day, both groups’ behaviour was tested in OFT and EPM. After that, animals had seven days of recovery and the injection and E-Stim session was repeated again. To verify if the non-phase-specific side effects of electrical stimulation had any influence on the OFT and EPM performance, the implanted group received a third round LPS i.p. with AntiPhase gamma E-Stim (Fig. 4A).

### Behavioural tests

#### Sucrose preference test

As mentioned above, all animals were housed separately after surgery in their homecage with four transparent plexiglas walls (rat chamber cage size: 38 cm width ×45 cm length × 18 cm height, plexiglas walls were 50 cm tall; mouse chamber with lid, cage size: 14×40×10 cm, plexiglas walls were 30 cm tall). Two holes were drilled through one of the walls in a certain distance from each other at the same height (rat chamber, 22 cm apart; mouse chamber, 6 cm apart). Two bottles with water pipes, one filled with tap water and the other with 1% (wt/vol) sucrose solution, were inserted through the holes under 15° angle to prevent leakage but giving free access to both liquids. For mouse experiments, the first day was for habituation, and liquid consumptions of subsequent days with daily alternating bottle locations were recorded for further analysis. In the rat 24 h SPT experiments, the bottles locations were also swapped every day and fresh sucrose solution and tap water were provided, unless otherwise noted. The daily consumption was recorded at 7 pm every day. The ratio of consumed sucrose solution relative to the total intake (water + sucrose solution) during 24 h is considered as the sucrose consumption (%) (*73*).

#### Open field test

The open field apparatus consisted a square arena (100×100 cm for rats, 50×50 cm for mice) with walls 40 cm height that was made of wooden board with sticky wallpaper (black or custom painted for rats, white for mice). The wooden floor was painted by light grey colour with a matte surface. The video camera was placed a certain distance on the top of apparatus. Before test, animals were transferred to the room for at least one hour habituation. Then animals were put into the center initially and recorded for 10 min. After each trial, the test zone was cleaned by 80% ethanol. The arena was divided into the center area (25% square corresponding to each apparatus for mice and rats respectively in the center) and the peripheral region. Body center was captured by EthoVision XT software (RRID: SCR_000441, Noldus, Wageningen, Netherlands), and the total distance travelled, time spent in the center area and number of entries into center in 10 min were automatically calculated (*74, 75*).

#### Elevated plus maze test

The maze was made of wooden board with a light grey painted matte floor, and consisted of two open arms and two closed arms (40 cm high walls with black wallpaper). Each arm was 50 cm long and 10 cm wide. The maze was standing on legs 50 cm above the floor of the room. The same camera and light intensity as in the OFT were used here. Rats were placed into center headed to the open arms, and video monitored for 10 mins. The time spent in the open arms, closed arms, center area and the total distance travelled during the 10 mins were quantified by EthoVision XT software (*76*).

#### Spontaneous movements

In the closed-loop gamma E-Stim experiments, the spontaneous movements of rats were continuously monitored by a camera mounted above the home cage from 7 to 9 pm during the SPT experiments.

### Histology

For verifying the virus expression and recording electrode placements, the animals were deeply anesthetized (1.5 g/kg urethane (i.p.)) and transcardially perfused with physiological saline (0.9% NaCl) followed by 4% paraformaldehyde (PFA) solution. For the implanted animals, one recording site in each brain area was lesioned with anodal direct current for 10 s before perfusion (Rat, 100 μA; mouse, 40 μA) (Fig. S7 A). Brains were post-fixed overnight in 4% PFA, sectioned to 50 μm thick slices using a vibrating microtome (VT1000S, Leica, Buffalo Grove, IL, USA). The slices were then stained with 1 μg/ml 4′, 6-Diamidino-2-phenylindole dihydrochloride (D8417; Sigma-Aldrich, St. Louis, MO, USA) in distilled water. Fluorescent signals were examined with a Zeiss LSM880 laser scanning confocal microscopy (Carl Zeiss, Oberkochen, Germany). Images were acquired using a Plan-Apochromat 20×/0.8 M27 or an alpha Plan-Apochromat 63×/1.46 Oil Korr M27 objective lens as previously described (*70*).

### Analysis

All LFP data analysis and statistical analysis were performed in MATLAB (RRID: SCR_001622; Mathworks, Natick, MA, USA). Behavioural analysis was performed using EthoVision XT software (RRID: SCR_000441, Noldus), unless otherwise stated.

#### Power spectrum and coherence analysis

Signals were pre-processed with down sampling to 1250 Hz if sampling rate was original 20 kHz. Power spectra were calculated in MATLAB using Multitaper Spectral Estimation in the Chronux Toolbox (http://chronux.org/) (Fig. S1 A and G). Coherence spectra between OB and other brain areas was calculated by using coherency function, which was based on the Multitaper coherency method from Chronux toolbox as well (Fig. S1 B–D). For all the analysis described above 3 s sliding windows with a 50% overlap were used (*77*). To test the correlation between gamma power and sucrose consumption in the optogenetic experiments, gamma power change was defined as [(PirC averaged gamma power from 10 min after illumination – PirC averaged gamma power from 10 min before illumination)/ PirC averaged gamma power from 10 min before illumination] (Fig. 2 G and H).

#### Off-line analysis of gamma events

Off-line detection of the gamma activity was employed in the acute CNO mouse/rat experiments (Fig. 1E, Fig. S2) and closed loop gamma E-Stim experiments (Fig. 3F–H, Fig. S8). First, the LFP was band-pass filtered with a eighth order zero phase lag Butterworth filter at 30–80 Hz, and RMS power was calculated in 50 ms sliding windows. Outliers of pooled power values were removed to get the mean and standard deviation of power values as reference. Gamma bursts were detected where the windowed power values exceeded 3 times of standard deviation (S.D.) above the mean value for that particular frequency for at least three consecutive windows. The boundaries of each gamma events were determined where the power values fall below mean + 2 S.D. around the previously identified peaks. Detection accuracy was confirmed by subjective visual observation. Sleep didn’t affect gamma event detection because gamma power was strongly reduced in the both OB and PirC during sleep. For the analysis of gamma incidence in the closed loop gamma E-Stim experiment, awake periods were classified first from 2 h home cage recordings.

#### Statistical analysis

Data are presented as mean ± S.D. Statistical testing was performed using MATLAB. Two-way repeated ANOVA followed by Tukey’s post hoc test was employed to compare behavioural performance in the chronic mice CNO experiment between the mCherry and the hM4Di groups. Wilcoxon rank-sum test was employed to compare gamma power changes between the two groups in the acute CNO mouse/rat experiments. Pearson’s correlation coefficient was used for testing the correlation between performance of SPT and reduction of gamma oscillations in the PirC and OB, respectively. One-way ANOVA test followed by Tukey’s post hoc test was employed to examine SPT among 24 h, WD + 2 h and WD + light +2 h in the optogenetic experiments, and among three phase-dependent electrical stimulus in the closed-loop gamma E-Stim experiments and also for gamma incidence in the latter experiments. Unpaired *t*-test was employed for power changes of gamma events. In the LPS experiments, unpaired *t*-test was employed to test SPT performance, Wilcoxon rank-sum test was used for testing the significance of other behavioural tests such as OFT and EPM. The significance level was set at *P* < 0.05. *, ** and *** indicate differences of *P* < 0.05, *P* < 0.01 and *P* < 0.001, respectively. For details of statistical analyses see Table S5.

## Supplementary figures

**Fig. S1.**
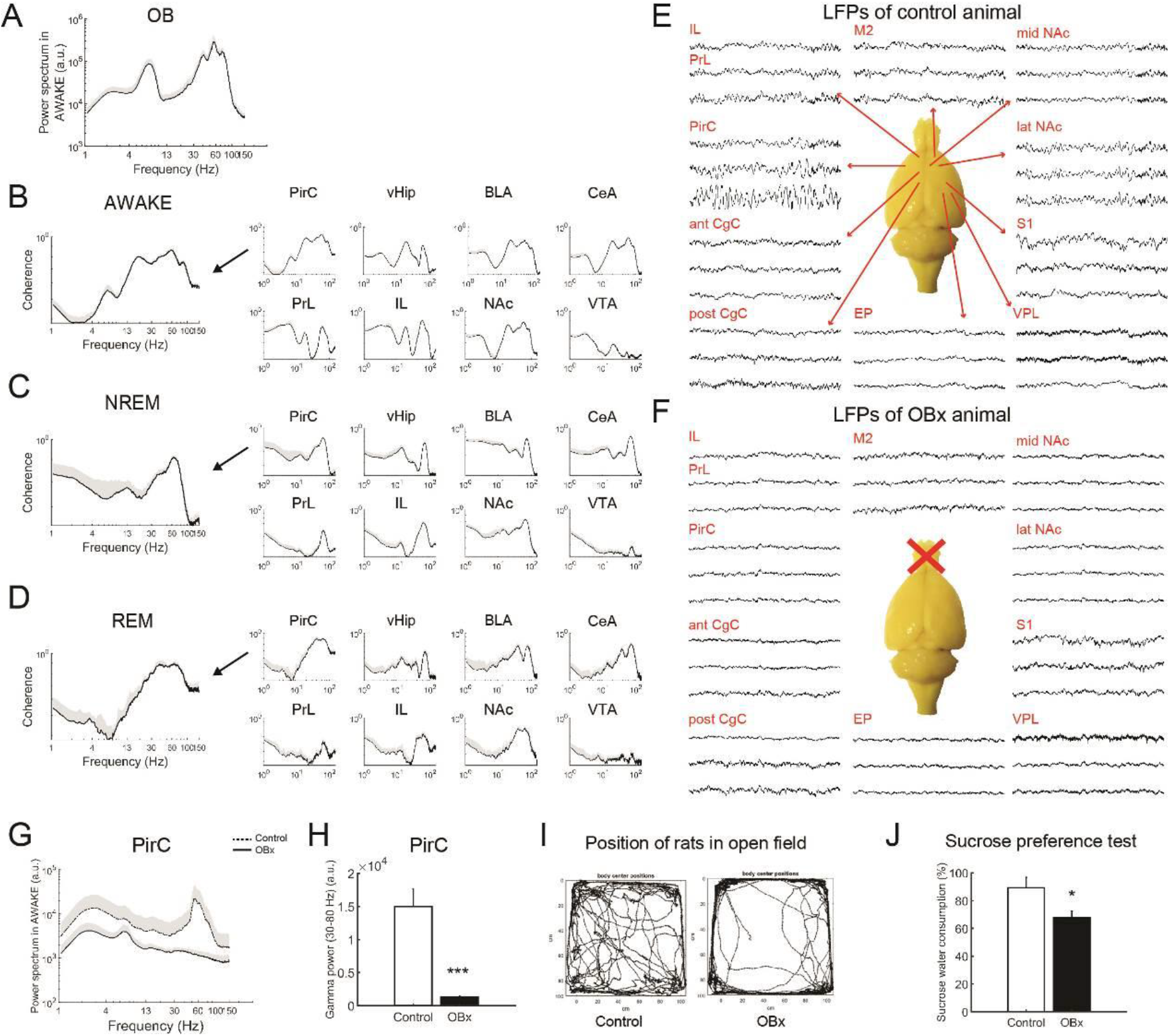
Olfactory bulbectomy reduces global gamma oscillations and induces depressive-like behaviors in rats. (**A**) Power spectrum of OB LFP in intact animals during AWAKE state. OB, olfactory bulb. (**B–D**) Highly coherent gamma oscillations between OB and multiple brain regions during AWAKE (B), NREM (C) and REM (D) states, respectively. PirC, piriform cortex; vHip, ventral hippocampus; CeA/BLA, central amygdala/basolateral amygdala; PrL/IL, prelimbic cortex/ infralimbic cortex; VTA, ventral tegmental area. (**E, F**) Representative local field potentials (LFPs) of control rat (E) and OBx rat (F) in multiple brain regions. M2, secondary motor cortex; mid NAc, medial nucleus accumbens; lat NAc, lateral nucleus accumbens; ant CgC, anterior cingulate cortex; post CgC, posterior cingulate cortex; S1, primary somatosensory cortex; EP, entopeduncular nucleus; VPL, ventral posterolateral thalamic nucleus. (**G**) Power spectrum in the PirC of control animal (dashed line) and OBx animal (bold line) during the AWAKE state. Grey shadow indicates S.D. (**H**) Statistical results in gamma band corresponding to (G). (**I**) Representative traces of control and OBx animals’ position in the open field in 10 mins one month after OBx. (**J**) Percentage of sucrose water consumption in both of groups one month after the surgery. (n = 3 rats/ group). Values are represented as means + S.D. *** indicates *P* < 0.0001.

**Fig. S2.**
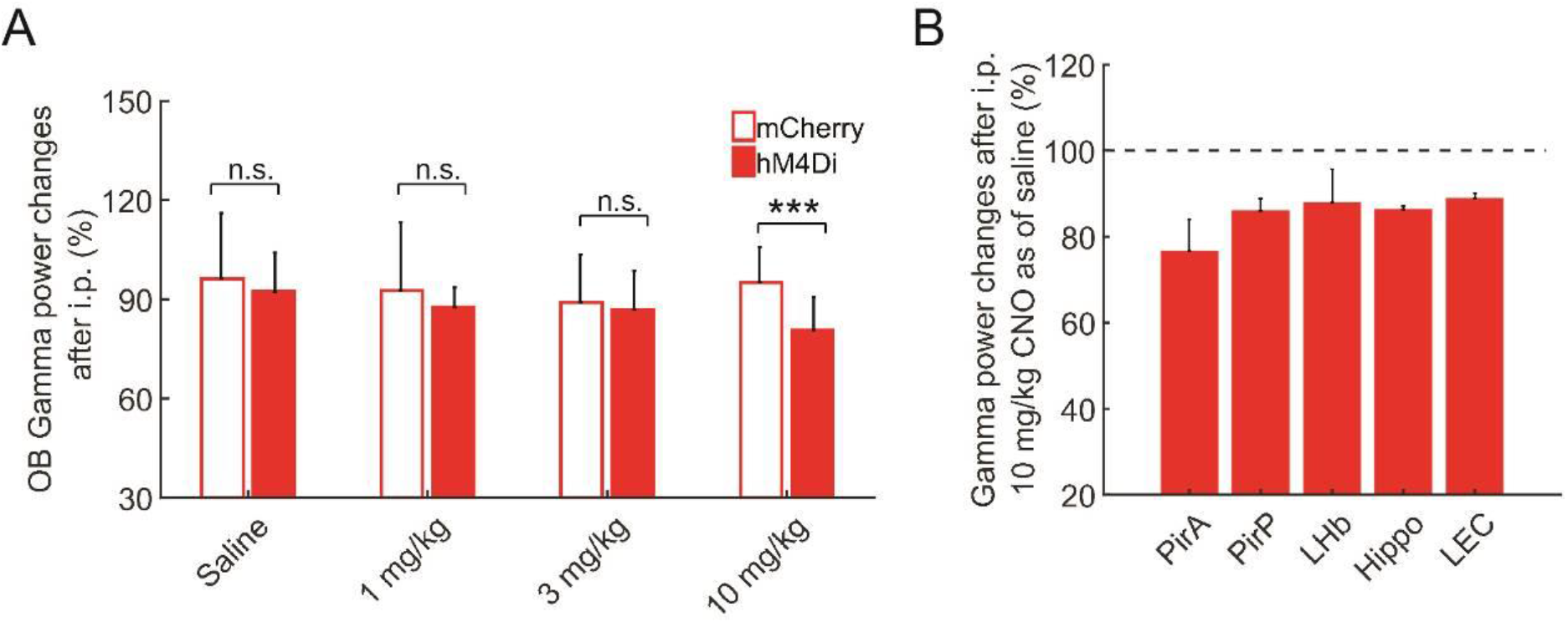
Chemogenetic inhibition of OB neurons decreases OB gamma oscillations in a dose dependent manner in rats. Related to Fig. 1. (**A**) Changes of OB gamma oscillations (30–80 Hz) after systemic administration of either saline or CNO in both of mCherry and hM4Di-mCherry groups. The protocol is the same as the four days long acute CNO experiments on mice (Fig. 1D). (**B**) Representative brain wide power changes of gamma oscillations after systemic administration of 10 mg/kg CNO in hM4Di rat. Values are represented as means + S.D. n.s., not significant; \*\*\**P* < 0.05.

**Fig. S3.**
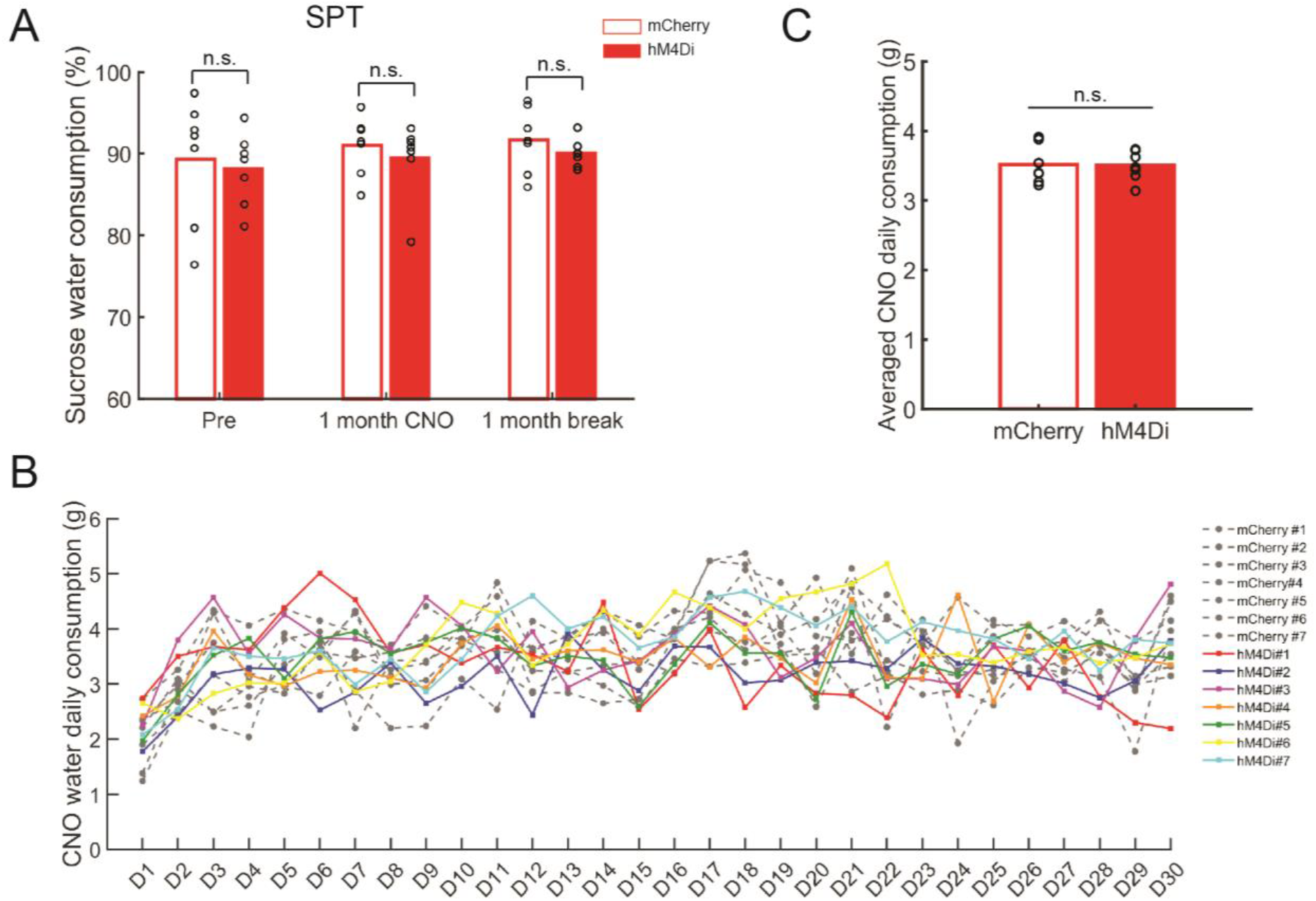
Chemogenetic inhibition of OB neurons doesn’t change the consumption of either CNO water or sucrose water. Related to Fig. 1. (**A**) No significant differences were found between the CNO treated hM4Di and control groups in the sucrose preference test (SPT) (n = 7 mice / group). (**B**) Individual daily CNO solution consumption of the animals. Each line represents one mouse. (**C**) No significant differences were found in the averaged CNO daily consumption (n = 7 mice / group). Circles and bars denote per animal averages and means across animals, respectively. . n.s., not significant.

**Fig. S4.**
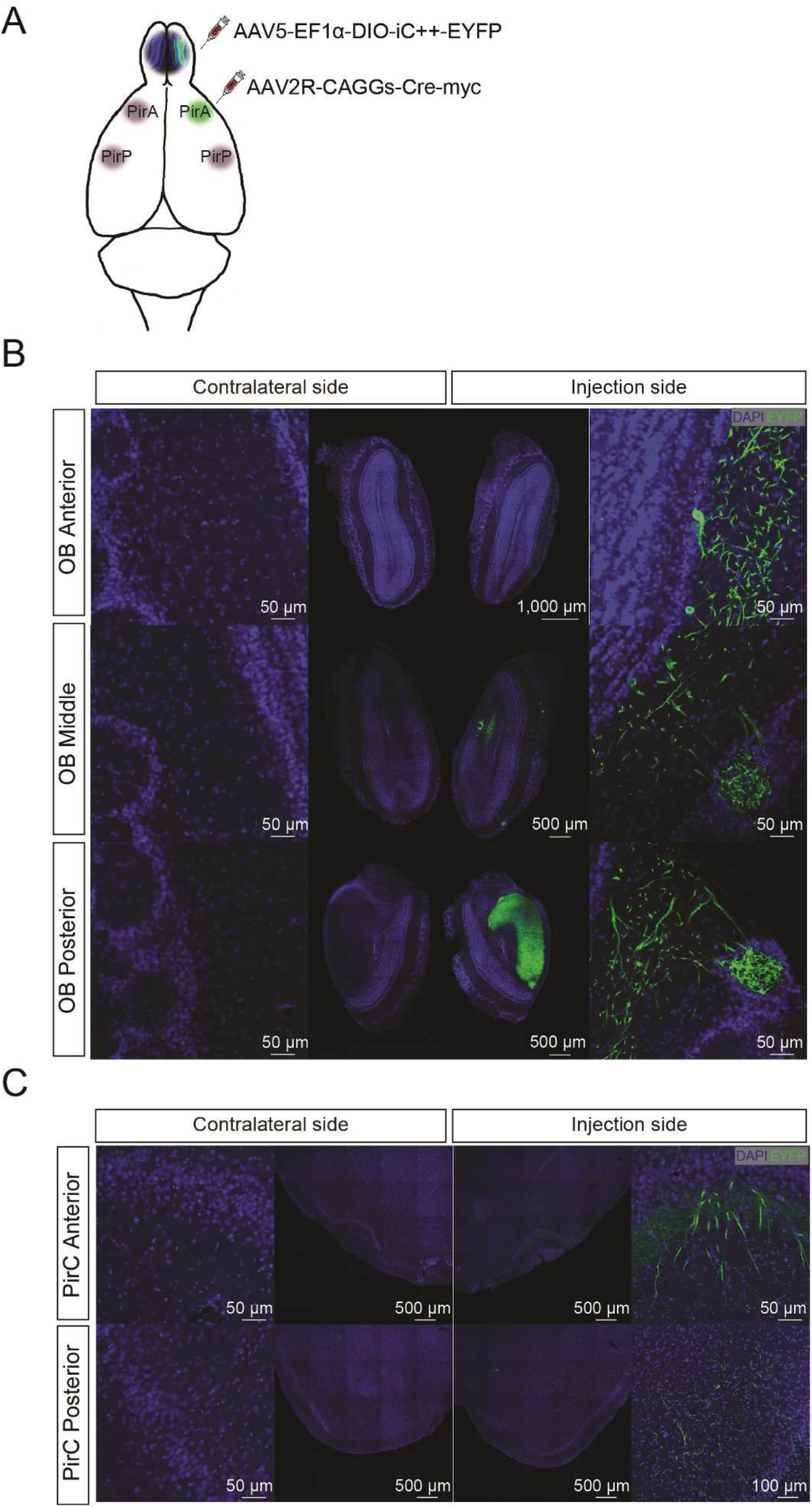
Visualization of OB neurons projecting to the anterior part of the ipsilateral PirC. Related to Fig. 2. (**A**) The schema of viral vector injections. (**B, C**) EYFP expression is present ipsilateral to the injection sites in the OB (**B**) and PirC (**C**), respectively, but not contralaterally.

**Fig. S5.**
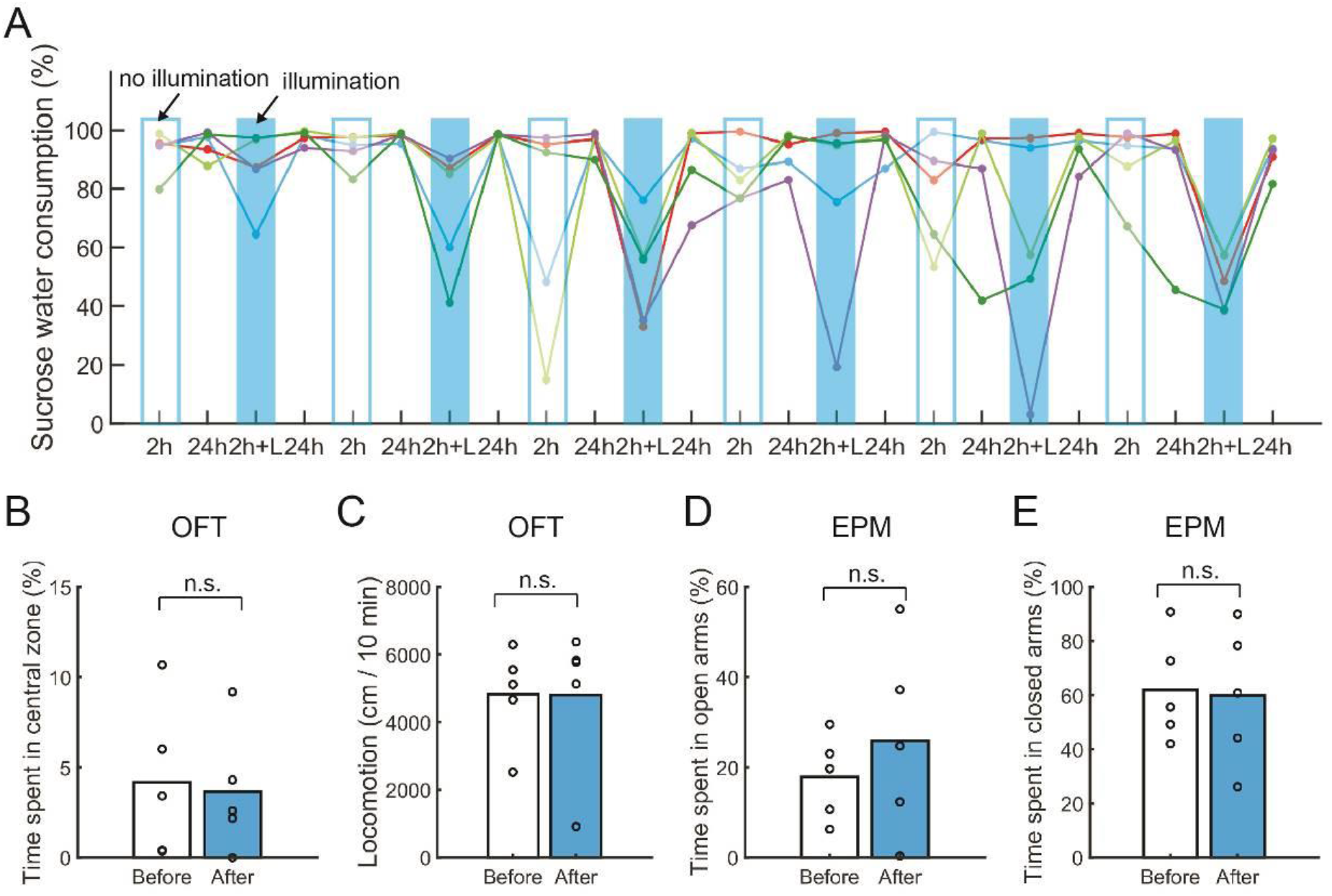
Behavioral performances during and after the selective, reversible suspension of synaptic transmission of OB to PirC pathway by optogenetic CALI (InSynC). Related to Fig. 2. (**A**) Time courses of sucrose preference of individual rats during the whole protocol as showed in Fig. 2E. Colored lines and markers indicate individual rats (n = 5). Open blue and solid blue bars mark test sessions with and without illuminations, respectively. 2 h, two hour SPT test after 22 hours water deprivation; 24 h, 24 hours SPT test without water deprivation; L, light/optostimulation. (**B, C**) No significant differences were found either in the time spent in the central zone (**B**) or locomotion (**C**) in the open field test (OFT), comparing before illumination (Before) and after illumination (After) measurements. (**D, E**) No significant differences were found either in the time spent of open arms (**D**) or closed arms (**E**) in the elevated plus maze (EPM) comparing before illumination and after illumination periods(n = 5). Circles and bars denote per animal and across animal averages, respectively. n.s., not significant.

**Fig. S6.**
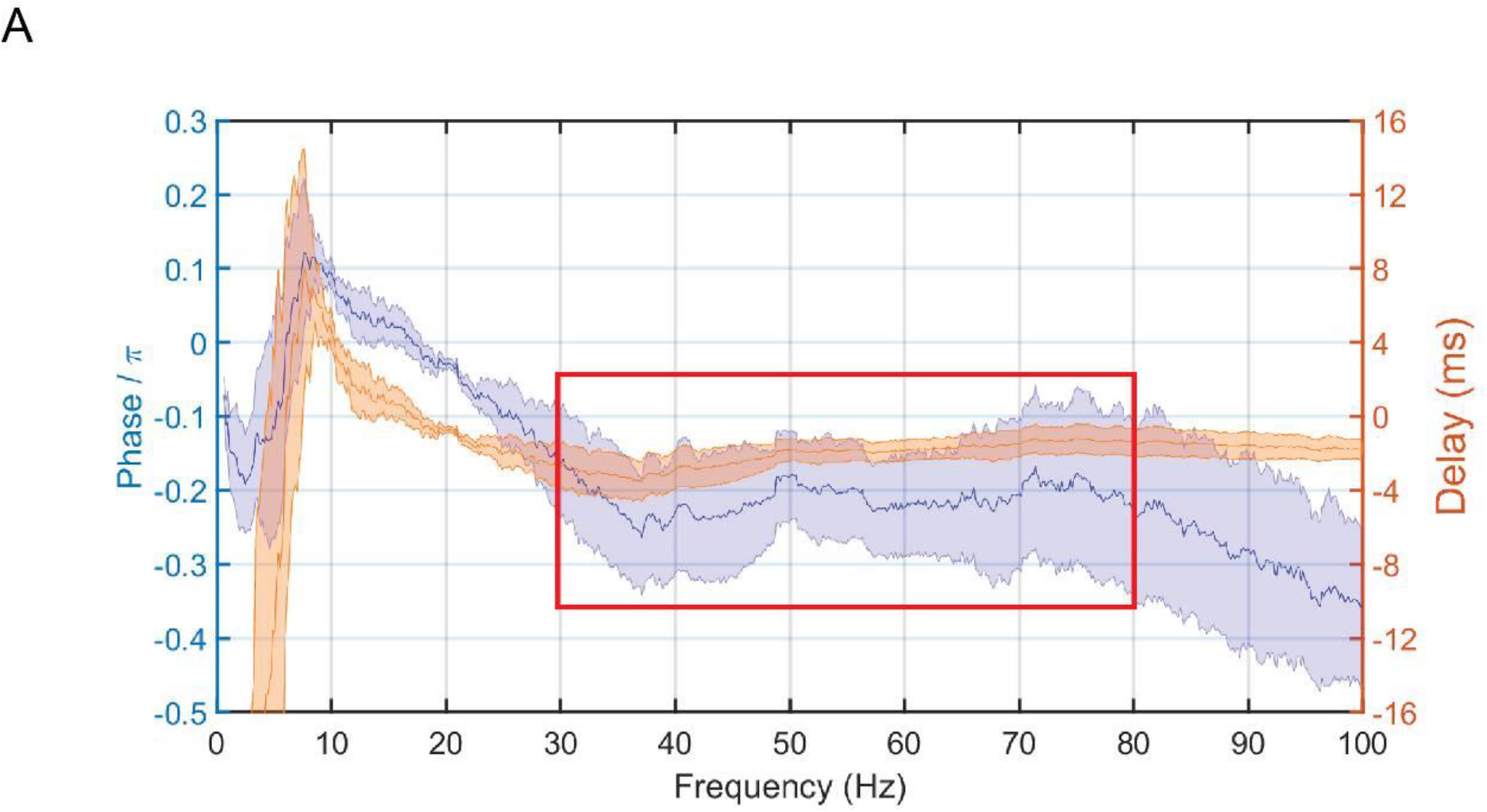
Phase lag coherency between OB and PirC in naïve rats during AWAKE state. Related to Fig. 3. (**A**) Phase lag coherency between OB and PirC from 0–100 Hz with a window width of 5 s during 10–30 min AWAKE state. The gamma band (30–80 Hz) of PirC signal is lagging OB activity with a - 0.21π ± 0.08 π. (n = 5 rats).

**Fig. S7.**
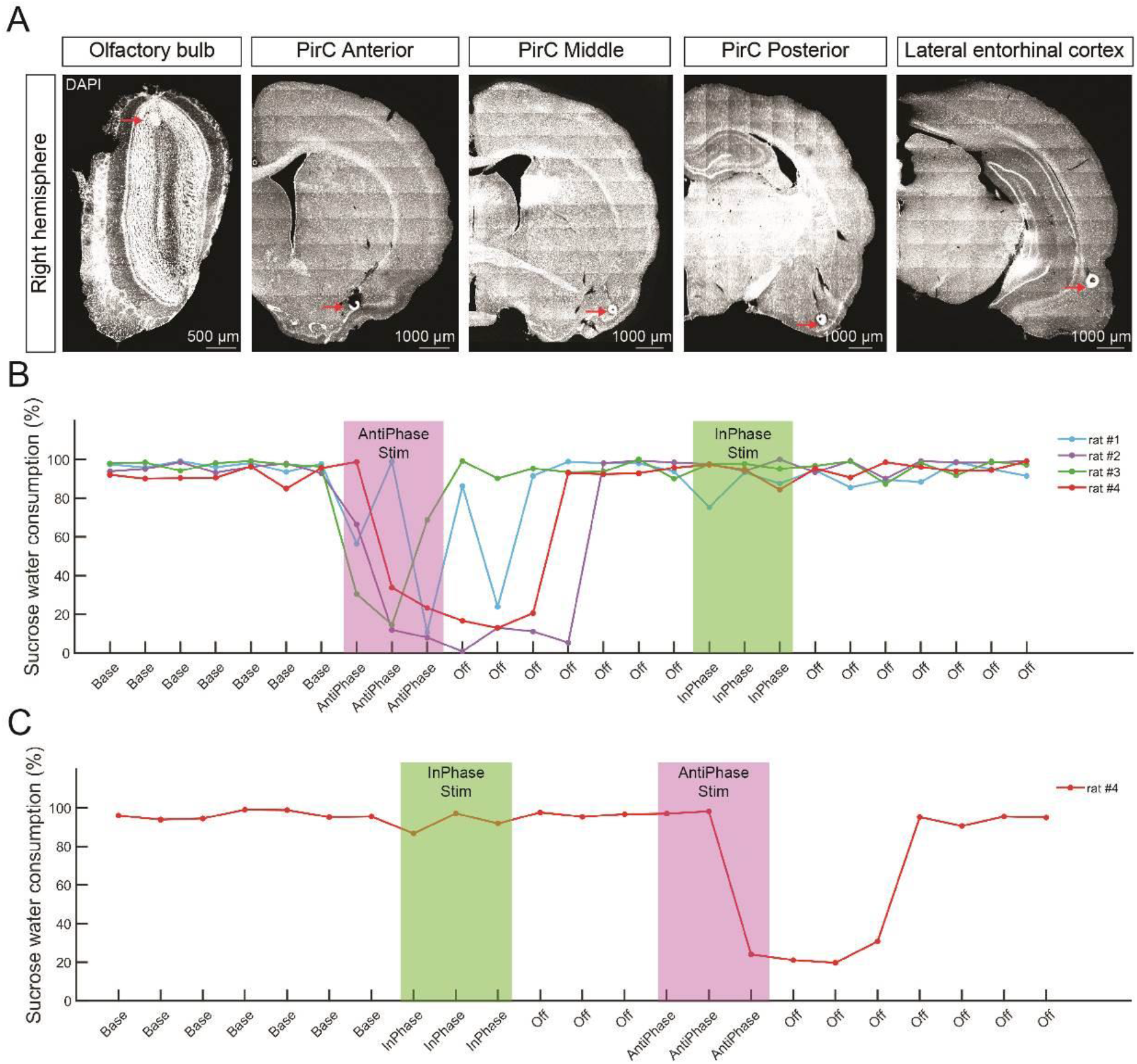
Time courses of sucrose water consumption of individual rats with real-time close-loop feed of OB gamma oscillations to the PirC. Related to Fig. 3. (**A**) Post-mortem identification of recording sites’ locations. Each arrow indicates a recording site. (**B**) Time course of sucrose water consumption of the four rats that underwent the AntiPhase - InPhase protocol as shown in Fig. 3C. Colored lines denote individual rats. (**C**) Time course of sucrose water consumption of the fourth rat that underwent the flipped sequence of InPhase - AntiPhase stimulation. The order of stimulation paradigms did not affect their performance.

**Fig. S8.**
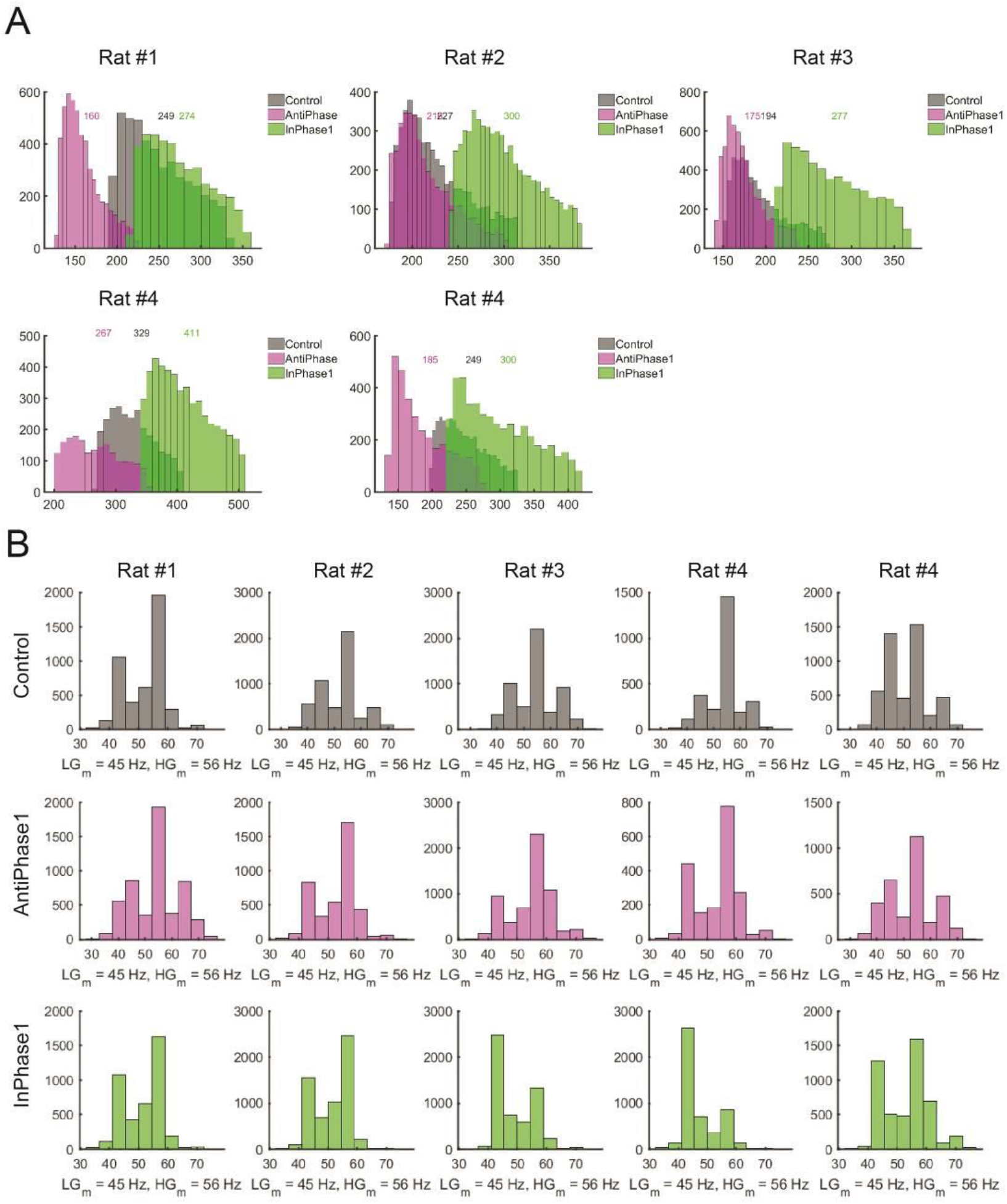
Features of gamma events in the PirC after real-time closed-loop feed of the OB gamma oscillations to PirC. Related to Fig. 3. (**A**) The power distribution of gamma events in each individual trial during one hour LFP recording during Baseline (grey), during theday after AntiPhase stimulation (magenta) and the day after InPhase stimulation (green). The numbers in each figure represent medians of the distributions. The conventions are the same as Fig. 3F. (**B**) The frequency distributions of gamma events of individual trials shown in (A). LG_m_ represents the median of frequency from 30 to 50 Hz, and HG_m_ represents the medians of frequency from 50 to 80 Hz. Five trials from four rats are as shown in Fig S6.

**Fig. S9.**
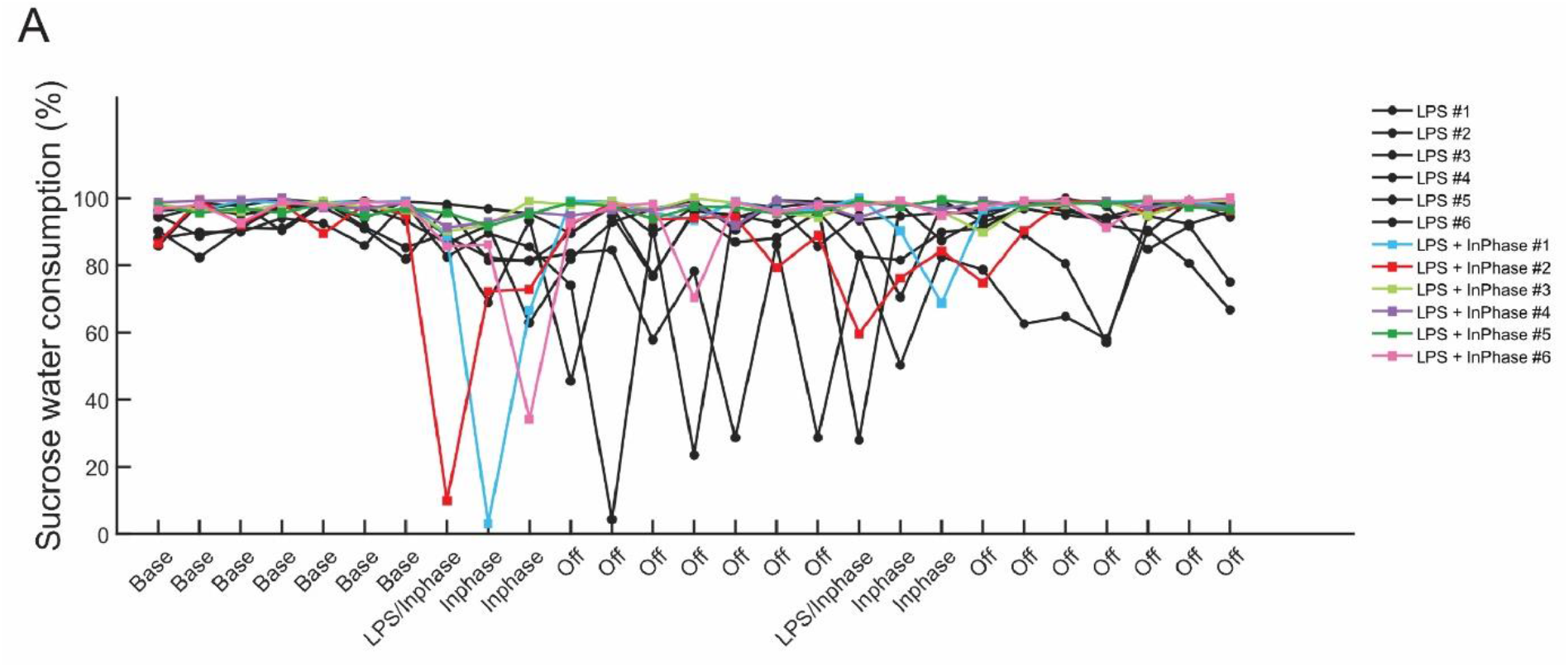
Time courses of sucrose water consumption during InPhase stimulation. Related to Fig. 4. (A) Time courses of sucrose water consumption of individual rats following two sessions of systemic LPS administrations. Related to the Fig. 4B group data. (n = 6 rats per group)

## Supplementary Tables

**Table S1.**
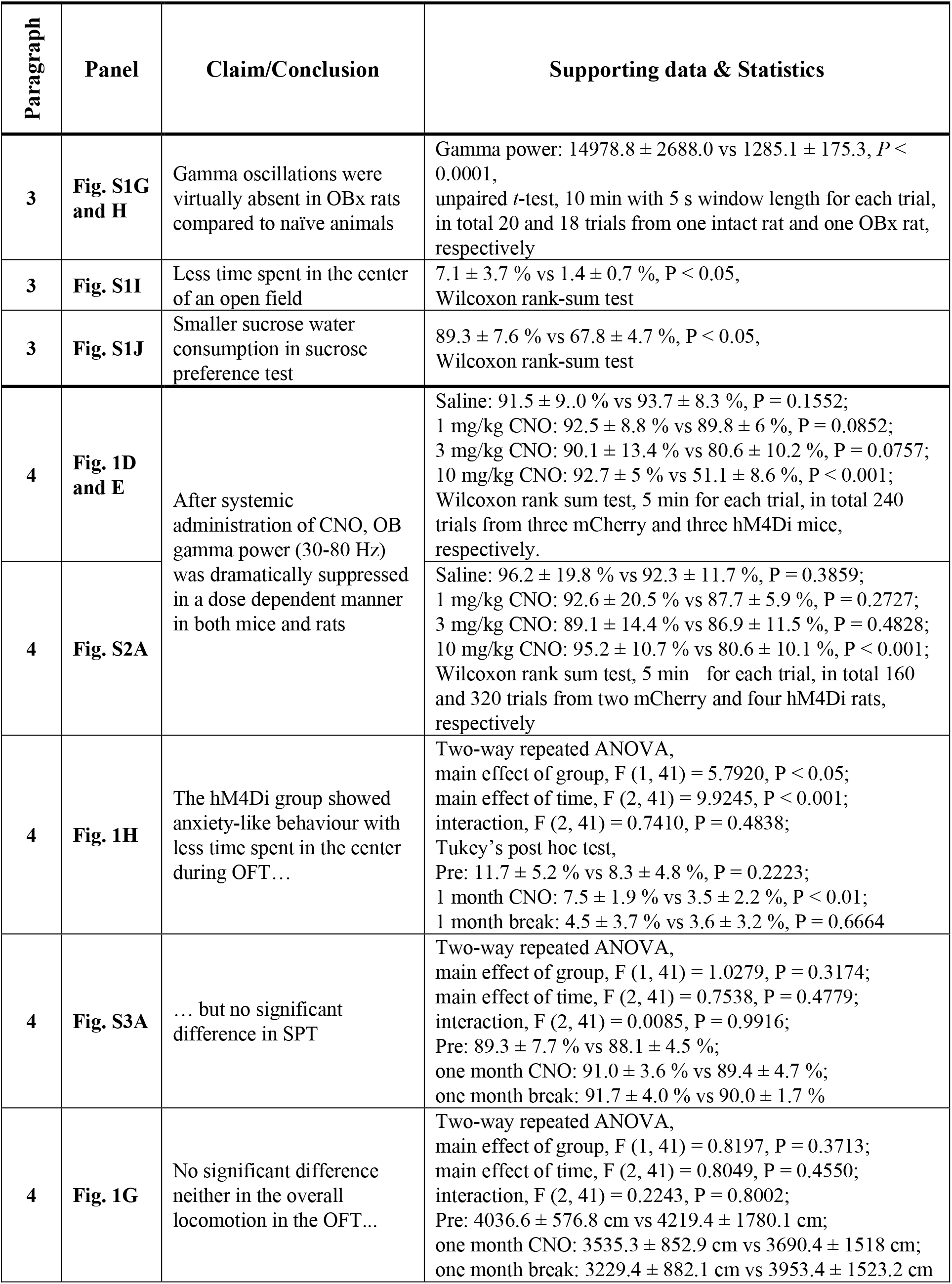
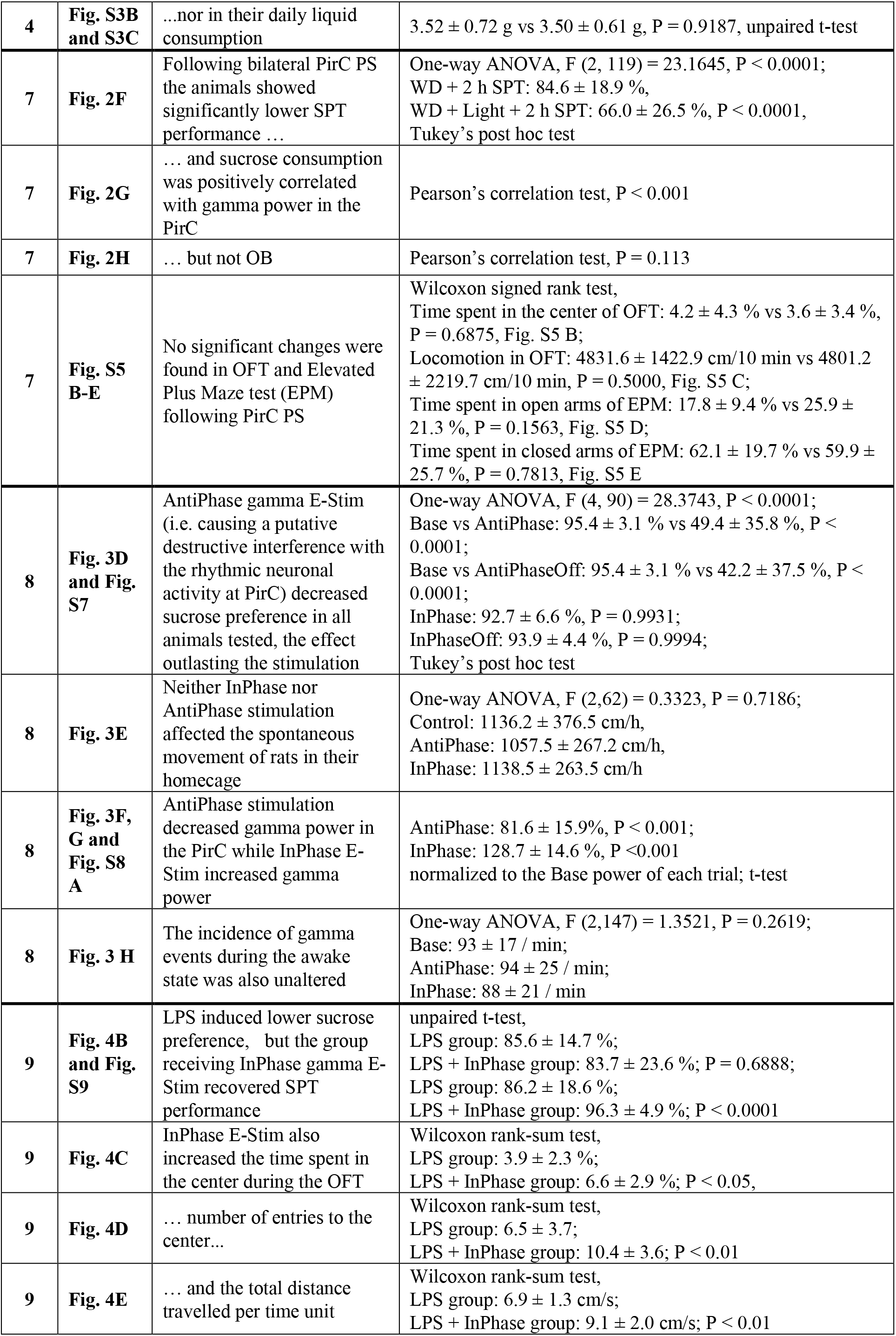
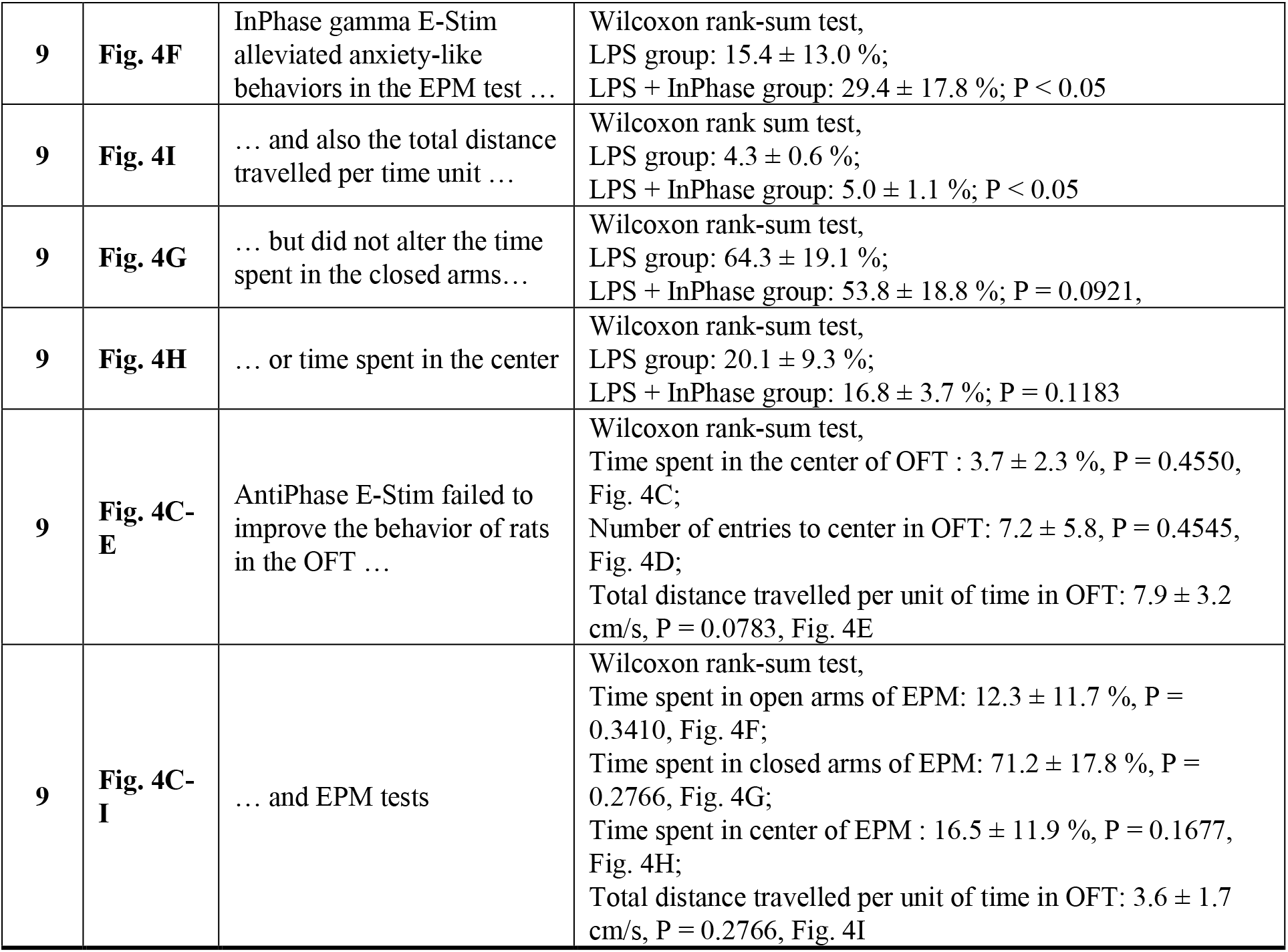
Results description table.

**Table S2.**
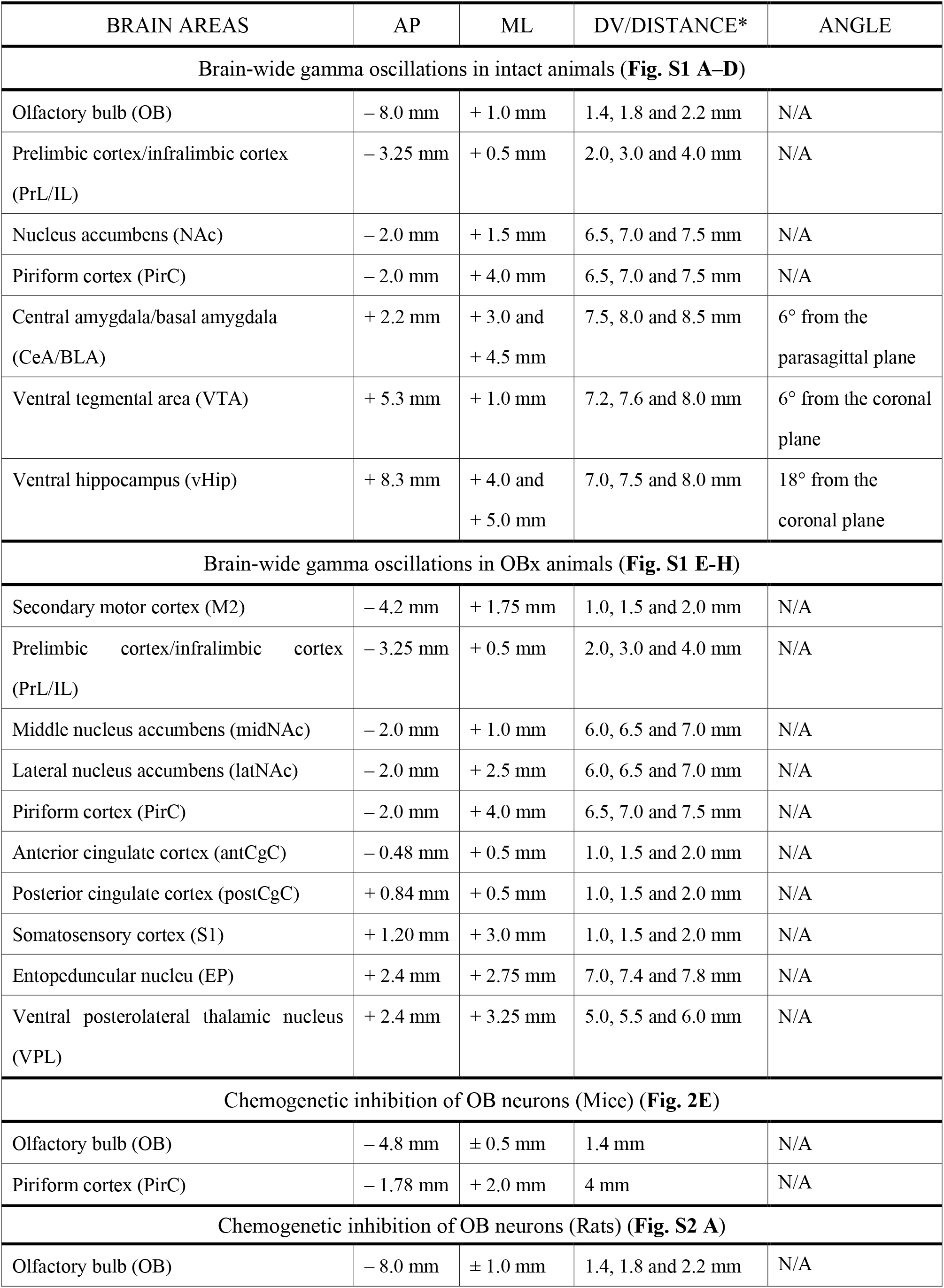
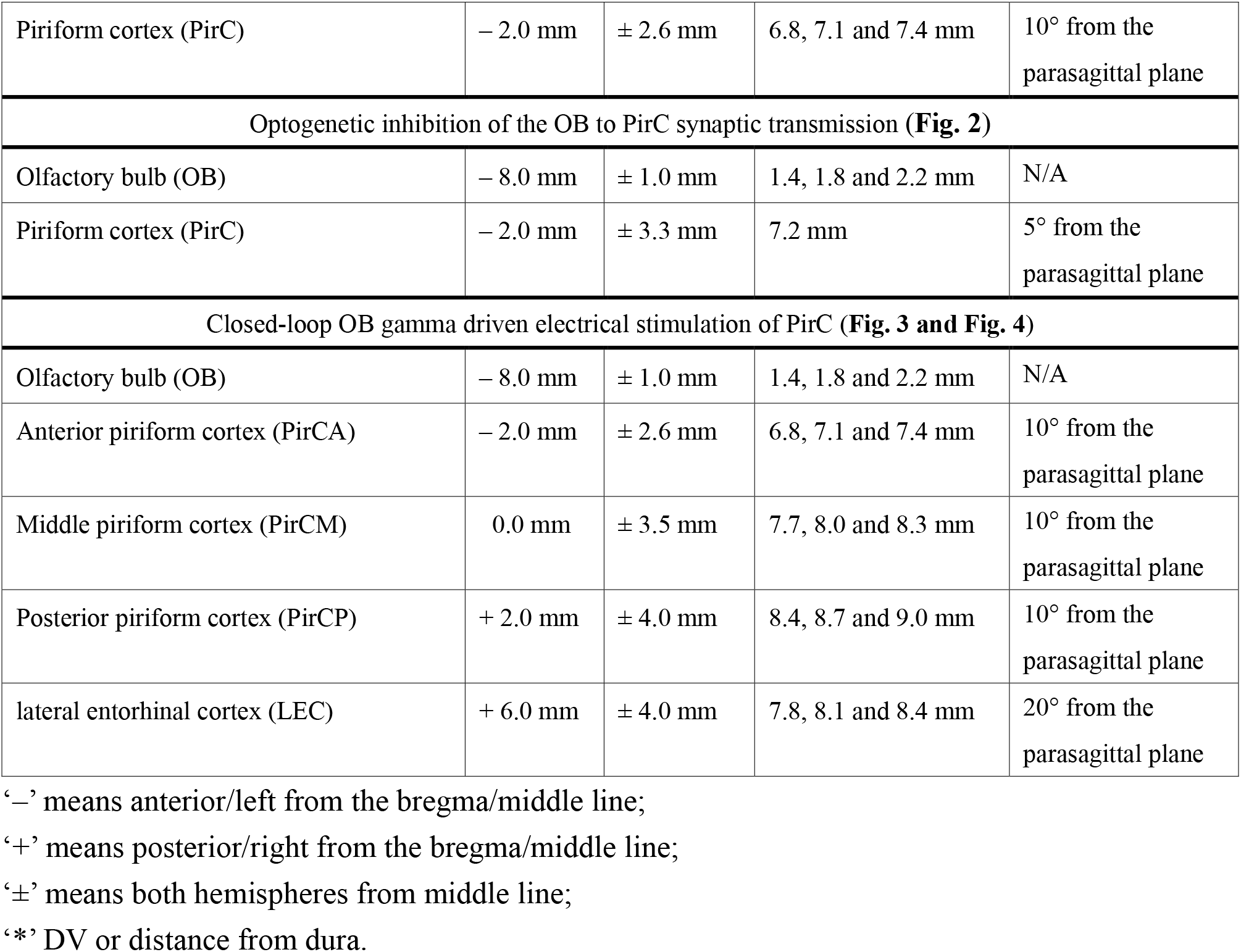
Electrodes implantation coordinates table.

| BRAIN AREAS | AP | ML | DV/DISTANCE* | ANGLE |
| --- | --- | --- | --- | --- |
| Brain-wide gamma oscillations in intact animals (**Fig. S1 A–D**) |  |  |  |  |
| Olfactory bulb (OB) | – 8.0 mm | + 1.0 mm | 1.4, 1.8 and 2.2 mm | N/A |
| Prelimbic cortex/infralimbic cortex (PrL/IL) | – 3.25 mm | + 0.5 mm | 2.0, 3.0 and 4.0 mm | N/A |
| Nucleus accumbens (NAc) | – 2.0 mm | + 1.5 mm | 6.5, 7.0 and 7.5 mm | N/A |
| Piriform cortex (PirC) | – 2.0 mm | + 4.0 mm | 6.5, 7.0 and 7.5 mm | N/A |
| Central amygdala/basal amygdala (CeA/BLA) | + 2.2 mm | + 3.0 and<br>+ 4.5 mm | 7.5, 8.0 and 8.5 mm | 6° from the parasagittal plane |
| Ventral tegmental area (VTA) | + 5.3 mm | + 1.0 mm | 7.2, 7.6 and 8.0 mm | 6° from the coronal plane |
| Ventral hippocampus (vHip) | + 8.3 mm | + 4.0 and<br>+ 5.0 mm | 7.0, 7.5 and 8.0 mm | 18° from the coronal plane |
| Brain-wide gamma oscillations in OBx animals (**Fig. S1 E–H**) |  |  |  |  |
| Secondary motor cortex (M2) | – 4.2 mm | + 1.75 mm | 1.0, 1.5 and 2.0 mm | N/A |
| Prelimbic cortex/infralimbic cortex (PrL/IL) | – 3.25 mm | + 0.5 mm | 2.0, 3.0 and 4.0 mm | N/A |
| Middle nucleus accumbens (midNAc) | – 2.0 mm | + 1.0 mm | 6.0, 6.5 and 7.0 mm | N/A |
| Lateral nucleus accumbens (latNAc) | – 2.0 mm | + 2.5 mm | 6.0, 6.5 and 7.0 mm | N/A |
| Piriform cortex (PirC) | – 2.0 mm | + 4.0 mm | 6.5, 7.0 and 7.5 mm | N/A |
| Anterior cingulate cortex (antCgC) | – 0.48 mm | + 0.5 mm | 1.0, 1.5 and 2.0 mm | N/A |
| Posterior cingulate cortex (postCgC) | + 0.84 mm | + 0.5 mm | 1.0, 1.5 and 2.0 mm | N/A |
| Somatosensory cortex (S1) | + 1.20 mm | + 3.0 mm | 1.0, 1.5 and 2.0 mm | N/A |
| Entopeduncular nucleu (EP) | + 2.4 mm | + 2.75 mm | 7.0, 7.4 and 7.8 mm | N/A |
| Ventral posterolateral thalamic nucleus (VPL) | + 2.4 mm | + 3.25 mm | 5.0, 5.5 and 6.0 mm | N/A |
| Chemogenetic inhibition of OB neurons (Mice) (**Fig. 2E**) |  |  |  |  |
| Olfactory bulb (OB) | – 4.8 mm | ± 0.5 mm | 1.4 mm | N/A |
| Piriform cortex (PirC) | – 1.78 mm | + 2.0 mm | 4 mm | N/A |
| Chemogenetic inhibition of OB neurons (Rats) (**Fig. S2 A**) |  |  |  |  |
| Olfactory bulb (OB) | – 8.0 mm | ± 1.0 mm | 1.4, 1.8 and 2.2 mm | N/A |
| Piriform cortex (PirC) | – 2.0 mm | ± 2.6 mm | 6.8, 7.1 and 7.4 mm | 10° from the parasagittal plane |
| Optogenetic inhibition of the OB to PirC synaptic transmission (**Fig. 2**) |  |  |  |  |
| Olfactory bulb (OB) | – 8.0 mm | ± 1.0 mm | 1.4, 1.8 and 2.2 mm | N/A |
| Piriform cortex (PirC) | – 2.0 mm | ± 3.3 mm | 7.2 mm | 5° from the parasagittal plane |
| Closed-loop OB gamma driven electrical stimulation of PirC (**Fig. 3 and Fig. 4**) |  |  |  |  |
| Olfactory bulb (OB) | – 8.0 mm | ± 1.0 mm | 1.4, 1.8 and 2.2 mm | N/A |
| Anterior piriform cortex (PirCA) | – 2.0 mm | ± 2.6 mm | 6.8, 7.1 and 7.4 mm | 10° from the parasagittal plane |
| Middle piriform cortex (PirCM) | 0.0 mm | ± 3.5 mm | 7.7, 8.0 and 8.3 mm | 10° from the parasagittal plane |
| Posterior piriform cortex (PirCP) | + 2.0 mm | ± 4.0 mm | 8.4, 8.7 and 9.0 mm | 10° from the parasagittal plane |
| lateral entorhinal cortex (LEC) | + 6.0 mm | ± 4.0 mm | 7.8, 8.1 and 8.4 mm | 20° from the parasagittal plane |

‘–’ means anterior/left from the bregma/middle line;
‘+’ means posterior/right from the bregma/middle line;
‘±’ means both hemispheres from middle line;
‘\*’ DV or distance from dura.

**Table S3.**
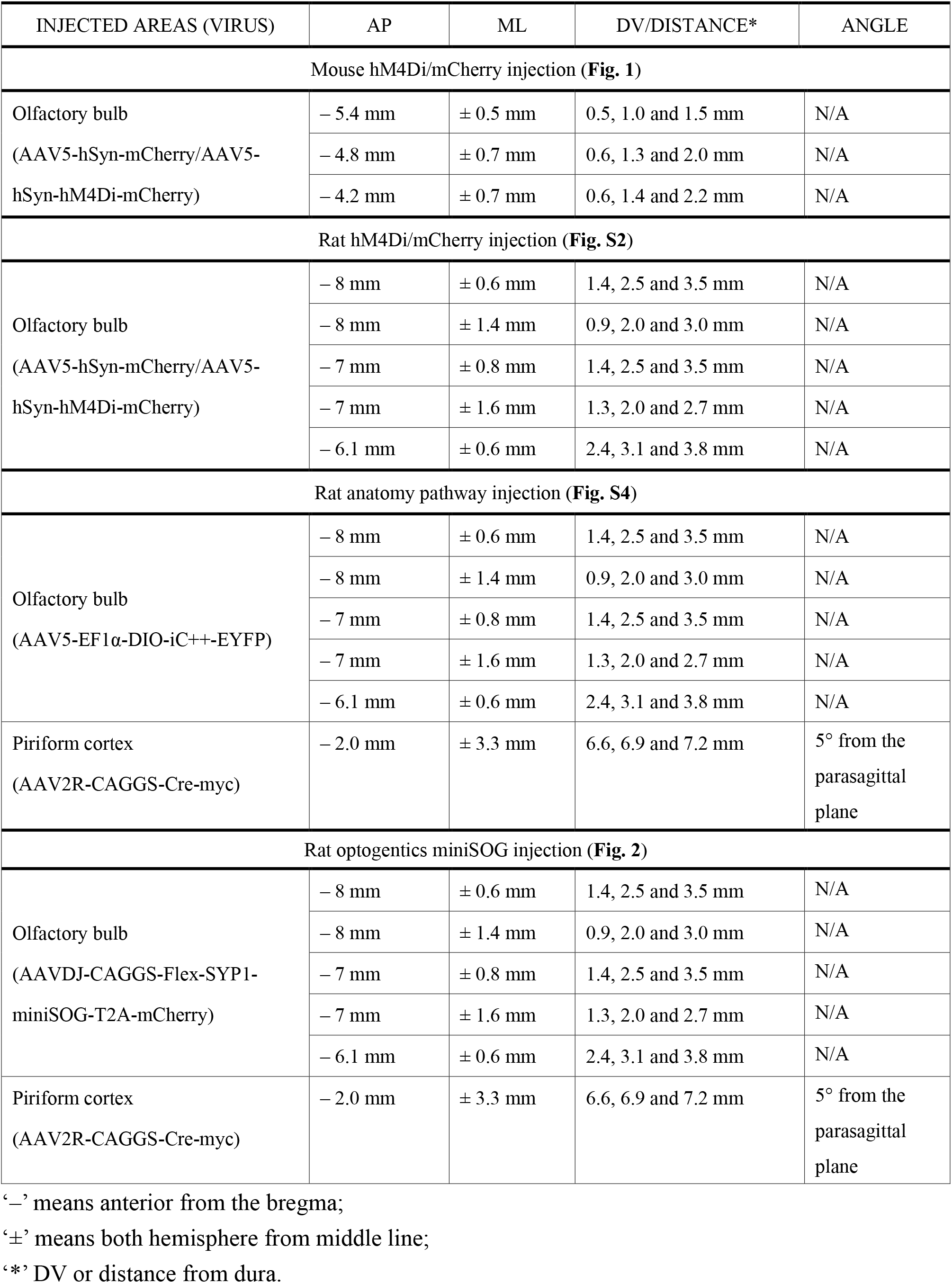
Virus injection coordinates table.

| INJECTED AREAS (VIRUS) | AP | ML | DV/DISTANCE* | ANGLE |
| --- | --- | --- | --- | --- |
| Mouse hM4Di/mCherry injection (**Fig. 1**) |  |  |  |  |
| Olfactory bulb<br>(AAV5-hSyn-mCherry/AAV5-hSyn-hM4Di-mCherry) | – 5.4 mm | ± 0.5 mm | 0.5, 1.0 and 1.5 mm | N/A |
|  | – 4.8 mm | ± 0.7 mm | 0.6, 1.3 and 2.0 mm | N/A |
|  | – 4.2 mm | ± 0.7 mm | 0.6, 1.4 and 2.2 mm | N/A |
| Rat hM4Di/mCherry injection (**Fig. S2**) |  |  |  |  |
| Olfactory bulb<br>(AAV5-hSyn-mCherry/AAV5-hSyn-hM4Di-mCherry) | – 8 mm | ± 0.6 mm | 1.4, 2.5 and 3.5 mm | N/A |
|  | – 8 mm | ± 1.4 mm | 0.9, 2.0 and 3.0 mm | N/A |
|  | – 7 mm | ± 0.8 mm | 1.4, 2.5 and 3.5 mm | N/A |
|  | – 7 mm | ± 1.6 mm | 1.3, 2.0 and 2.7 mm | N/A |
|  | – 6.1 mm | ± 0.6 mm | 2.4, 3.1 and 3.8 mm | N/A |
| Rat anatomy pathway injection (**Fig. S4**) |  |  |  |  |
| Olfactory bulb<br>(AAV5-EF1α-DIO-iC<sup>++</sup>-EYFP) | – 8 mm | ± 0.6 mm | 1.4, 2.5 and 3.5 mm | N/A |
|  | – 8 mm | ± 1.4 mm | 0.9, 2.0 and 3.0 mm | N/A |
|  | – 7 mm | ± 0.8 mm | 1.4, 2.5 and 3.5 mm | N/A |
|  | – 7 mm | ± 1.6 mm | 1.3, 2.0 and 2.7 mm | N/A |
|  | – 6.1 mm | ± 0.6 mm | 2.4, 3.1 and 3.8 mm | N/A |
| Piriform cortex<br>(AAV2R-CAGGS-Cre-myc) | – 2.0 mm | ± 3.3 mm | 6.6, 6.9 and 7.2 mm | 5° from the parasagittal plane |
| Rat optogenetics miniSOG injection (**Fig. 2**) |  |  |  |  |
| Olfactory bulb<br>(AAVDJ-CAGGS-Flex-SYP1-miniSOG-T2A-mCherry) | – 8 mm | ± 0.6 mm | 1.4, 2.5 and 3.5 mm | N/A |
|  | – 8 mm | ± 1.4 mm | 0.9, 2.0 and 3.0 mm | N/A |
|  | – 7 mm | ± 0.8 mm | 1.4, 2.5 and 3.5 mm | N/A |
|  | – 7 mm | ± 1.6 mm | 1.3, 2.0 and 2.7 mm | N/A |
|  | – 6.1 mm | ± 0.6 mm | 2.4, 3.1 and 3.8 mm | N/A |
| Piriform cortex<br>(AAV2R-CAGGS-Cre-myc) | – 2.0 mm | ± 3.3 mm | 6.6, 6.9 and 7.2 mm | 5° from the parasagittal plane |

**Table S4.** Key Resource Table.

| REAGENT or RESOURCE | SOURCE | IDENTIFIER |
| --- | --- | --- |
| Bacterial and Virus Strains |  |  |
| AAV5-hSyn-mCherry | Addgene | Cat# 114472-AAV5 |
| AAV5-hSyn-hM4Di-mCherry | Addgene | Cat# 50475-AAV5 |
| pAAV-hSyn-mCherry | Addgene | Cat# 14772 |
| pAAV-hSyn-hM4D(Gi)-mCherry | Addgene | Cat# 50475 |
| pAAV-SYP1-miniSOG-T2A-mCherry | Addgene | Cat# 50972 |
| AAVDJ-CAGGS-Flex-SYP1-miniSOG-T2A-mCherry | This paper | N/A |
| AAV2R-CAGGS-Cre-myc | This paper | N/A |
| AAV5-EF1α-DIO-iC<sup>++</sup>-EYFP | UNC Vector Core | N/A |
| Chemicals, Peptides, and Recombinant Proteins |  |  |
| Clozapine N-oxide | Sigma-Aldrich | Cat# C0832 |
| Lipopolysaccharides from Escherichia coli O111:B4 | Sigma-Aldrich | Cat# L2630 |
| Urethane | Sigma-Aldrich | Cat# U2500 |
| Paraformaldehyde | Sigma-Aldrich | Cat# P6148 |
| DAPI | Sigma-Aldrich | Cat# D8417 |
| Software and Algorithms |  |  |
| MATLAB R2017b | Mathworks | RRID:SCR_001622 |
| Neuroscope | Hazan et al., 2006<br>(14)(14) | RRID:SCR_002455 |
| EthoVision XT software | Noldus | RRID:SCR_000441 |
| ZEN Digital Imaging for Light Microscopy | Carl Zeiss | RRID:SCR_013672 |
| Circular Statistics Toolbox | Berens et al., 2009<br>(15)(15) | RRID:SCR_016651 |
| Other |  |  |
| HML Insulated Tungsten 99.95% wire | California Fine Wire | Cat# CFW2044436 |

**Table S5. Statistical table**

Provided as a separate Excel file.

**Table S6. Mouse CNO water daily consumption**

Provided as a separate Excel file.

